# Pyramidal cell types drive functionally distinct cortical activity patterns during decision-making

**DOI:** 10.1101/2021.09.27.461599

**Authors:** Simon Musall, Xiaonan R. Sun, Hemanth Mohan, Xu An, Steven Gluf, Shujing Li, Rhonda Drewes, Emma Cravo, Irene Lenzi, Chaoqun Yin, Björn M. Kampa, Anne K. Churchland

**Affiliations:** Institute of Biological Information Processing (IBI-3), Forschungszentrum Jülich, Jülich, Germany; Department of Neurophysiology, Institute for Zoology, RWTH Aachen University, Aachen, Germany; Cold Spring Harbor Laboratory, Neuroscience, Cold Spring Harbor, NY, USA; Department of Neurosurgery, Zucker School of Medicine, Hofstra University, Hempstead, NY, USA; Department of Neurobiology, Duke University Medical Center, Durham, NC; Department of Neurobiology, David Geffen School of Medicine, University of California, Los Angeles, CA, USA; JARA Brain, Institute for Neuroscience and Medicine, Forschungszentrum Jülich, Germany

## Abstract

Understanding how cortical circuits generate complex behavior requires investigating the cell types that comprise them. Functional differences across pyramidal neuron (PyN) types have been observed within cortical areas, but it is not known whether these local differences extend throughout the cortex, nor whether additional differences emerge when larger-scale dynamics are considered. We used genetic and retrograde labeling to target pyramidal tract (PT), intratelencephalic (IT) and corticostriatal projection neurons and measured their cortex-wide activity. Each PyN type drove unique neural dynamics, both at the local and cortex-wide scale. Cortical activity and optogenetic inactivation during an auditory decision task also revealed distinct functional roles: all PyNs in parietal cortex were recruited during perception of the auditory stimulus, but, surprisingly, PT neurons had the largest causal role. In frontal cortex, all PyNs were required for accurate choices but showed distinct choice-tuning. Our results reveal that rich, cell-type-specific cortical dynamics shape perceptual decisions.

## Introduction

The neocortex is organized into discrete layers that form a vertically-arranged microcircuit motif. This core circuit is largely conserved across cortical areas with each layer consisting of distinct excitatory and inhibitory cell types that can be categorized based on genetic markers, cell morphology, anatomical projections or developmental lineage^1^. The precise interplay between these cell types is crucial for accurate cortical circuit function and their respective functional roles are the subject of intense study. Tremendous progress has been made particularly for cortical interneurons, where the availability of specific mouse driver lines has revealed the functional arrangement of inhibitory circuit motifs^2–4^, e.g. for network synchronization^5–7^ and state-dependent sensory processing^8–11^. However, the roles of glutamatergic pyramidal neuron (PyN) types are less well established, although PyNs comprise ∼80% of all cortical neurons and form almost all long-range projections that enable the communication between local cortical circuits and other brain areas.

While often treated as a monolithic group, PyNs appear to be far more diverse than interneurons with at least one hundred putative subtypes indicated by RNA sequencing^12^. These molecular signatures are critical for categorizing PyNs and go far beyond layer identity as different subtypes are often intermingled within the same layers^13–17^. PyNs can also be classified based on their projection target. Long-range projection neurons are broadly categorized into two major types: intratelencephalic (IT) neurons, projecting to other cortical structures and the striatum, and pyramidal tract (PT) neurons, projecting to subcortical structures, such as the superior colliculus (SC), thalamus, the pons and the striatum. Furthermore, PT and IT neurons differ in their electrophysiological properties, dendritic morphology, local connectivity, and responsiveness to sensory stimulation^15–17^. Recent studies in sensory cortex also showed that only PT but not IT neurons are important for active perception of tactile or visual stimuli, suggesting that PT and IT neurons encode separate streams of information^18,19^. Similar results have been found in secondary motor cortex (M2), where specific PT neurons are involved in motor generation^13,20^. This suggests that the functional divergence of PyN types could be key for understanding cortical microcircuits, with PT and IT neurons forming functionally-distinct, parallel subnetworks that independently process different information. However, the functional tuning of individual PyNs in frontal cortex is still best predicted by cortical area location instead of laminar location or projection type^21^. Since PyN subtype activity has only been studied in single areas, it is therefore not known whether PyN-specific subcircuits exist throughout the cortex or only in a subset of cortical regions.

An ideal method to address this question is widefield calcium imaging, which allows measuring neural activity across the dorsal cortex with cell-type specificity^22–24^. Interneuron-specific widefield imaging revealed clear differences in the spatiotemporal dynamics of different inhibitory cell types during an odor detection task^25^. However, cortex-wide studies of different PyN types are lacking, in part due to the limited availability of PyN-specific driver lines^26–28^. Here, we used two novel knock-in mouse driver lines targeting PT or IT neurons^26^ and performed widefield Ca^2+^ imaging to measure PyN subtype-specific activity while animals performed a perceptual decision-making task. Moreover, we developed a retrograde labeling approach to selectively measure the activity of corticostriatal projection (CStr) neurons throughout the dorsal cortex. Dimensionality-reduction and clustering analyses revealed unique cortex-wide dynamics for each PyN subtype, suggesting the existence of specialized subcircuits. Cortical dynamics of different PyNs were further segregated based on their role in decision-making, with encoder and decoder approaches revealing the strongest stimulus- and choice-related modulation in sensory, parietal and frontal cortices. This was confirmed by PyN-type-specific inactivation experiments. In parietal cortex, PT neurons were most important for sensory processing while all PyN types in frontal cortex were needed for choice formation and retention. Taken together, our results demonstrate that different PyN-types exhibit functionally distinct, cortex-wide neural dynamics with separate roles during perceptual decision-making.

## Results

To monitor PyN-type specific neural activity throughout the dorsal cortex, we used two knock-in inducible CreER lines that target developmentally-distinct classes of excitatory cortical neurons: Fezf2-CreER targeting PT neurons and PlexinD1-CreER targeting IT neurons^26^. For PyN-type specific GCaMP6s expression, both lines were crossed with the Ai162 reporter line^29^ and CreER activity was induced at four weeks of age. In each mouse line, GCaMP expression was restricted to specific cortical layers and PyN types (Fig 1a,b). In Fezf2 mice, expression was concentrated in layer 5b (Fig. 1a). We observed axonal projections to multiple subcortical regions, such as striatum and the corticospinal tract, confirming that Fezf2 is a reliable marker for corticofugal PT neurons. In PlexinD1 mice, GCaMP expression was restricted to layer 2/3 and layer 5a and axonal projections were visible in the corpus callosum and the striatum (Fig. 1b). These results are in agreement with earlier reports^26^, confirming that PlexinD1 is a reliable marker for intracortical and corticostriatal IT neurons.

**Figure 1.**
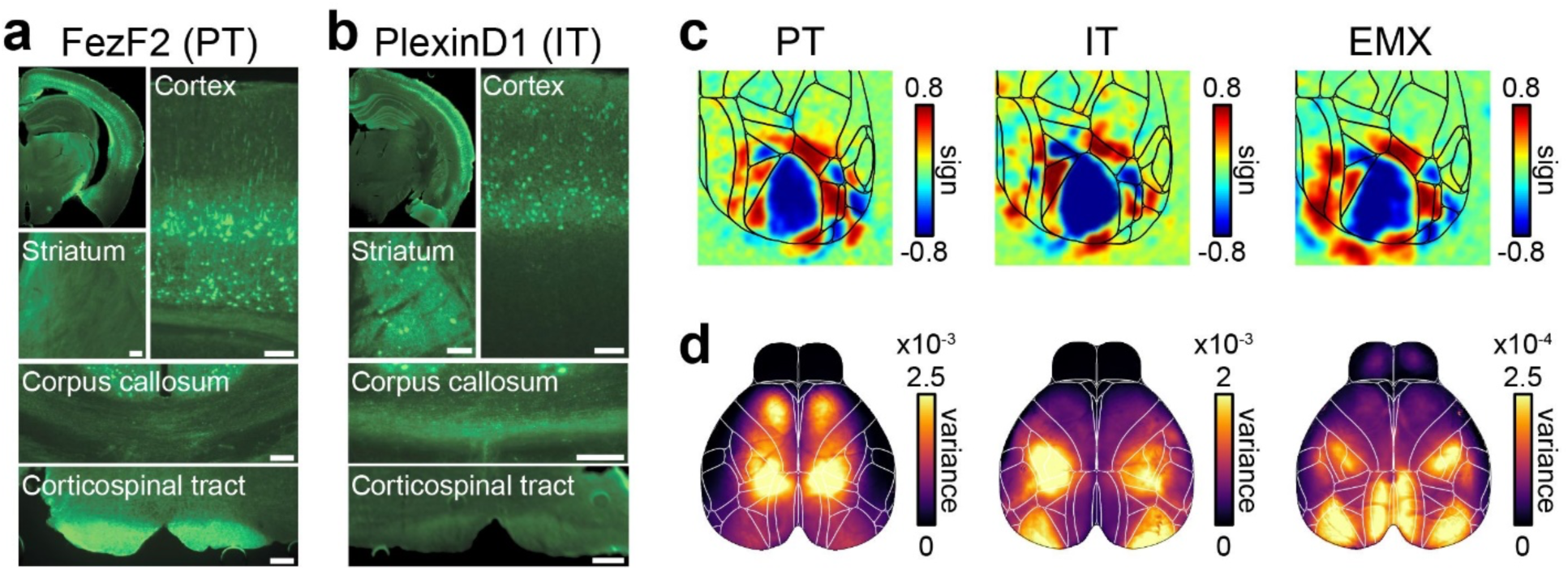
Knock-in mouse lines enable PyN subtype-specific recordings of cortex-wide neural activity. **a)** GCaMP6s expression in Fezf2-CreER;Ai162 mice. Cortical labeling is largely confined to layer 5b. Axonal projections were found in multiple subcortical regions such as the striatum and the corticospinal tract. (Scale bars: 100 µm). **b)** GCaMP6s expression in PlexinD1-CreER; Ai162 mice is widespread throughout the cortex and restricted to superficial layers 2/3 and layer 5a. Axonal projections were found in striatum and corpus callosum but absent in the corticospinal tract. **c)** Visual sign maps from retinotopic mapping experiments. IT and PT populations showed clear retinotopic responses in primary and secondary visual areas where boundaries largely resembled known areas that were also observed in nonspecific PyNs (EMX mice). **d)** Total variance maps from same mice as in c), showing most modulated cortical regions in each PyN type.

After confirming non-overlapping expression patterns of PT and IT neurons, we measured cell-type-specific cortical activity with widefield calcium imaging. All imaging data was aligned to the Allen Common Coordinate Framework v3 (CCF)^25,30^ to compare activity across individuals and PyN types. Both lines yielded robust GCaMP-dependent fluorescence and we observed rich neural dynamics throughout the dorsal cortex (Supp. Movies 1-3). We first used retinotopic mapping to assess sensory responses of different PyN types and the resulting spatial arrangement of visual areas. Both PT and IT neurons robustly responded to visual stimulation and we could construct retinotopic maps that reliably indicated the location of known visual areas (Fig. 1c)^31^. Retinotopic maps were similar to those observed in an Ai93D;Emx-Cre;LSL-tTA mouse line (EMX) that expressed GCaMP6f in all excitatory cortical neurons^32^, suggesting that the functional architecture of visual areas is comparable across PyN types. However, clear differences were apparent in the modulation of cortical regions in the absence of visual stimulation (Fig. 1d, same individual mice as in 1c). For example, the total variance of cortex-wide activity was largest in parietal and frontal regions in PT neurons (Fig. 1d, left) while variance was highest in visual and somatosensory regions of IT neurons (Fig. 1d, center). This was comparable to variance in EMX mice, which showed additional modulation in retrosplenial (RS) cortex (Fig. 1d, right). These different patterns were also highly consistent across individual mice of the same PyN type (Supp. Fig. 1).

The observed differences in variance could indicate general differences in the spatiotemporal dynamics of cortex-wide activity across PyN types. To describe such differences, we performed semi-nonnegative matrix factorization (sNMF), reducing the imaging data to a small number of spatial- and temporal components that capture nearly all variance (>99%)^23,33^. Here, dimensionality (the number of required components) is a measure of the overall complexity of cortical dynamics. Surprisingly, PT neurons had much lower dimensionality compared to IT neurons (Fig. 2a), suggesting that PT neurons are more functionally-homogeneous. This was also seen in the correlations between cortical regions, which were consistently highest for PT neurons (Supp. Fig. 2). A potential reason for this difference could be that IT neurons encompass a larger number of specialized subtypes than PT neurons and may thus support a wider range of functions^34^.

**Figure 2.**
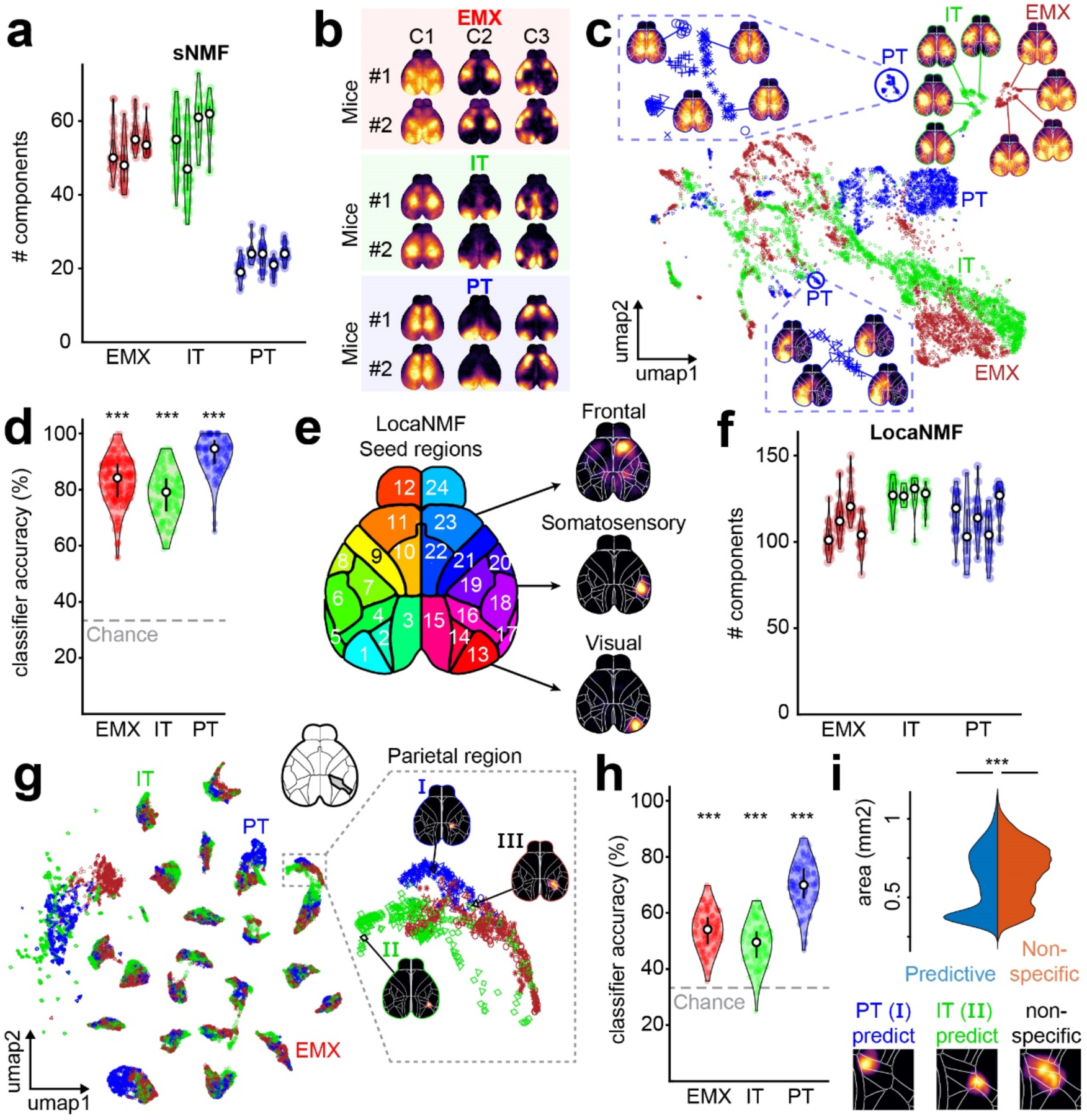
PyN subtypes exhibit unique cortical activity patterns. **a)** Number of sNMF components accounting for 99% of cortical variance in each PyN type. Violin plots show individual mice, dots represent individual sessions. **b)** Examples of spatial sNMF components. Components from different mice from the same PyN subtype (colored rectangles) strongly resembled each other. **c)** UMAP embedding of spatial sNMF components for EMX (red), IT (green) and PT (blue) mice. Maps show example spatial components for each type and their respective UMAP locations. Different markers denote individual mice. Blow-ups show examples of PT-specific regions. Additional clustering for individual mice is sometimes evident (top left region), but was generally weaker than PyN-specific clustering. **d)** Accuracy of a PyN type classifier based on neighbor identity of individual components in UMAP space. Each data point represents the mean classification accuracy over all components in one session. **e)** Map of seed regions used for LocaNMF analysis. Example spatial components are compact and mostly confined to the appropriate seed region. **f)** Number of LocaNMF components, accounting for 99% of cortical variance in each PyN type. Conventions as in (a). **g)** UMAP projection embedding of spatial LocaNMF components. Conventions as in (c). Left: UMAP shows clustering of LocaNMF components from similar regions (same 24 regions as shown in e). Right: Components within individual regions are further divided for different PyN types. Maps show example LocaNMF components (I-III). **h)** Accuracy of a PyN type classifier, based on individual LocaNMF components. Conventions as in (d). **i)** Top: Peak normalized distributions of area size for PyN-type-predictive (blue) versus unspecific (red) LocaNMF components. PyN-predictive components are smaller than unspecific components (PyN-predictive: median = 0.59 mm^2^, n = 6317 components, nonspecific: median = 0.68mm^2^, n = 18938 components; ranksum test: p < 10^−10^). Bottom: Examples of PyN-predictive (I, II) and unspecific (III) components in right parietal cortex.

Since PT neurons are a subset of EMX neurons, their lower dimensionality could indicate that PT components are also a subset of EMX components. We therefore quantified to which extent components from different PyN types are linearly overlapping or independent from each other. For each PyN type, we computed the amount of variance that could be explained with components from other mice of either the same or different PyN type (Supp. Fig. 3a). For all groups, components from a similar PyN type explained significantly more variance (ΔR^2^) than the maximal amount that could be explained by other types (EMX, mean ΔR^2^ = 3.3%, *t*-test: *p* < 10^−10^; IT, mean ΔR^2^ = 1.5%, *p* < 10^−10^; PT, mean ΔR^2^ = 1.2%, *p* < 10^−10^). Each group therefore contains additional PyN-type-specific variance that is independent from other types. This was also visible in reconstructed data: PT components captured larger fluctuations in IT activity (Supp. Movie 4) but failed to represent spatial details in the cortical activity patterns (Supp. Fig. 3b). Despite the similar dimensionality of IT and EMX activity (Fig. 2a), EMX ΔR^2^ was also similar for IT and PT components (ΔR^2^_EMX-IT_ = 3.4%, ΔR^2^_EMX-PT_ = 3.3%, *p* = 0.1), demonstrating that EMX data was not largely dominated by either IT or PT neurons.

To directly compare spatial activity patterns, we then focused on spatial sNMF components. Spatial components represent cortex-wide maps of positively correlated areas and we wondered whether these correlation patterns would differ between PyN subtypes. Indeed, individual spatial components from different mice of the same PyN subtype often strongly resembled each other but differed from other PyN subtypes (Fig. 2b). To assess if most spatial components (or only a small subset) was PyN-type-specific, we performed a UMAP projection of the first 20 components from all recordings and PyN types, non-linearly embedding the pixels of each component in a 2-dimensional space (Fig. 2c)^35^. If neural activity in PT, IT and EMX mice tended to exhibit similar correlation patterns, these spatial components would be mixed together. Instead, in agreement with the notion that components of the same PyN type resembled each other, we found that components formed clusters that were largely dominated by either PT or IT neurons (green/blue markers). EMX neurons formed a third set of non-overlapping clusters, likely reflecting the combined cortical dynamics from diverse PyN types beyond PT and IT neurons that were contained in this larger group (red markers). A simple classifier reliably identified each group, based on the nearest neighbors in a UMAP projection that was exclusively based on data from other animals. Remarkably, prediction accuracy was high even when the classification was based on just a single spatial component (Fig. 2d). The presence of such clear clusters is striking given that the spatial components used in the analysis were pooled over many sessions and mice. This confirms that UMAP clusters reflect consistent PyN-type-specific activity patterns, rather than idiosyncratic differences within individual sessions or mice. Taken together, these results clearly demonstrate that PyN types differ in the complexity of cortical dynamics, contain independent variance, and exhibit unique cortex-wide correlation patterns.

An important concern is that variations in the density of Cre expression in each mouse line could result in non-uniform GCaMP expression patterns which might contribute to PyN-specific spatial components. However, GCaMP-related fluorescence was largely uniform across cortex in all mice and we observed no clear relationship between raw widefield fluorescence patterns and PyN-specific spatial components (Supp. Fig. 4a, b). Moreover, despite some fluctuations in the density of Cre-expressing neurons across areas in both mouse lines, Cre-expression patterns did not account for spatial components or specific differences between PyN types (Supp. Fig. 4c, d). Nevertheless, particularly distinct activity of each PyN type in specific cortical areas could lead to cortex-wide correlation patterns that are either dominated by highly active areas or where inactive areas are ‘missing’. In this case, the relatively low dimensionality of PT neurons might be due to a lower number of active cortical areas. We therefore used a localized form of sNMF (LocaNMF)^33^ which extracts components that are dense and spatially restricted to a specific cortical “seed” region (Fig. 2e). Analyzing LocaNMF components therefore allowed us to reveal if PyN-specific differences mostly occur on a cortex-wide level (reflecting the interactions between cortical areas) or further extend to the local level of individual cortical areas (e.g. due to PyN-specific differences in the shape and localization of cortical areas).

First, we again compared the number of components required to explain at least 99% of the variance in each PyN type. As expected, the number of components was greater than sNMF and, interestingly, was also more similar across PyN subtypes (Fig. 2f). This shows that PyN-type-specific differences in cortex-wide correlation patterns are not due to a lack of activity in some cortical areas (which would have resulted in a lower number of required components, e.g. in PT mice) and instead result from PyN-type specific differences in the coordinated activation of multi-area cortical networks. Moreover, as with sNMF components, LocaNMF components from mice of the same PyN type explained more variance than components from other types (Supp. Fig. 3c).

Repeating the UMAP embedding for these local components also uncovered PyN-type-specific clustering (Fig. 2g), which could be used to accurately identify each PyN type (Fig. 2h). Classification accuracy was high across most cortical regions (Supp. Fig. 4d). Specificity of LocaNMF components could either be due to the presence of specific ‘subregions’, where PyN types are most active in smaller parts of a given cortical area, or larger ‘superregions’, where the activity of a specific PyN type extends across known area borders. We therefore compared the size of LocaNMF components that accurately predicted their respective PyN type (classifier accuracy > 99%) versus nonspecific components. Interestingly, PyN-predictive clusters were significantly smaller than nonspecific clusters (Fig. 2i), suggesting that different PyNs might be most active in distinct subregions instead of larger multi-area components. This indicates that smaller, PyN-specific subregions may reside within the coarser, traditionally-defined cortical areas.

We next assessed how cortical dynamics of PyN subpopulations are related to decision-making by imaging PyN subtype activity in animals trained on an auditory decision-making task (Fig. 3a)^36^. Mice initiated trials by touching small handles, which triggered the simultaneous presentation of sequences of clicking sounds to their left and right side. After a short delay period, mice reported decisions by licking one of two water spouts and were rewarded for choosing the side where more clicks were presented. To reduce temporal correlations between task events, such as trial initiation and stimulus onset, the durations of the initiation, stimulus and delay periods were randomly varied across trials. In all mice, decisions varied systematically with stimulus strength (Fig. 3b) and were equally affected by click sounds throughout the stimulus period (Supp. Fig. 5).

**Figure 3.**
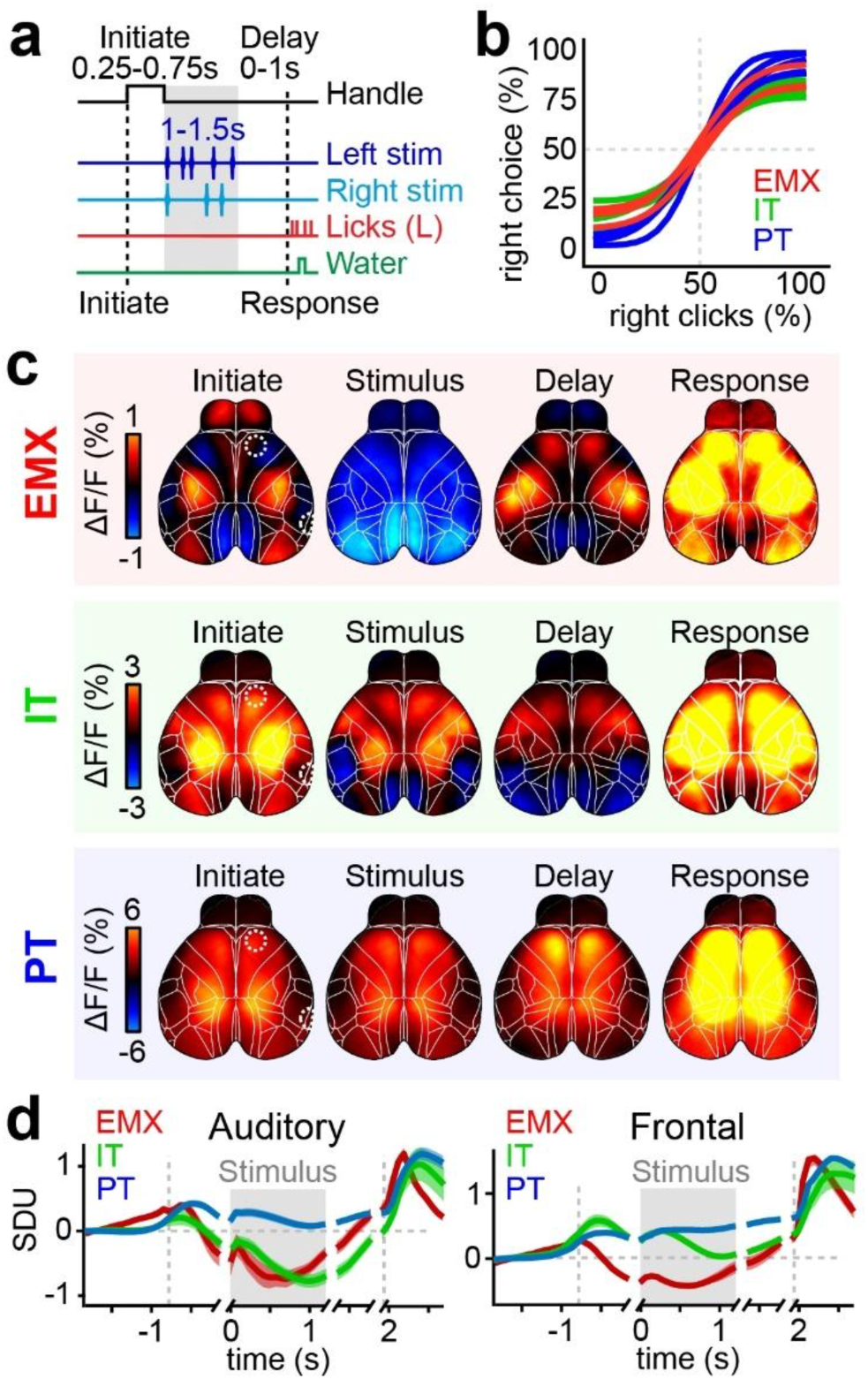
An auditory decision-making task reveals distinct functional activity patterns in each PyN type. **a)** Auditory discrimination task structure of an example trial. Mice touched paw handles to initiate randomized click sequences on the left and/or right side. After a delay period, a lick response on the correct side was rewarded with water. The episode duration was randomly varied in individual trials. **b)** Psychometric functions fit to behavioral data from the discrimination task in (a) of individual EMX (red), IT (green) and PT (blue) mice. **c)** Trial-averaged response maps for all correct, leftward trials in different PyN types. Shown are averages for the ‘Initiation’, ‘Stimulus’, ‘Delay’ and ‘Response’ periods shown in (a). **d)** Averaged activity in auditory (left) and frontal cortex (right) for each PyN type. Averages were separately aligned to each of the four trial periods, indicated by short gaps. Left dashed line: time of initiation, gray box: stimulus presentation, right dashed line: animal’s response. Traces show standard deviation units (SDU). White dashed circles in (c) show respective area locations. Colors as in (b). Shading shows s.e.m.; n = 4 EMX/IT mice and 5 PT mice.

As with spatial components, trial-averaged temporal sNMF and LocaNMF components showed pronounced clustering, suggesting that different PyN-types also exhibit distinct task-related temporal dynamics (Supp. Fig. 6). Correspondingly, we found clear differences in trial-averaged neural activity between PyN subtypes, especially during stimulus presentation (Fig. 3c). Here, cortical activity was uniformly suppressed in EMX mice, partially suppressed in somatosensory and visual areas in IT mice, and uniformly increased in PT mice (Fig. 3d). In all PyN types, activity was largely symmetric across the left and right hemispheres, even when only analyzing trials where stimuli were presented on the left side and the animal made a corresponding leftward choice (Fig. 3c; Supp. Fig. 7). Moreover, responses to sensory stimuli were much weaker than movement-related responses, such as trial initiation or licking (Fig. 3d, gray bar versus dashed lines). A potential explanation is that lateralized, task-related activity is obscured by cortical activity due to animal movements^32,37–39^.

To isolate task-related activity from movements we used a linear encoding model^32^. The model combines many task- and movement-variables to predict single-trial fluctuations in cortical activity (Fig. 4a, top). Task variables were related to the learned behavior, and included past and current choices, and the presentation of a sensory stimulus. Movement variables included licking and touching the handles, alongside facial movements (whisking or nose movements) obtained from two video cameras of the animal’s face and body (a complete list of all task- and movement variables is shown in Supp. Table 1). All variables were combined into a single design matrix and we used ridge regression to fit the model to the imaging data. This produced a time-varying event kernel for each variable, allowing us to isolate how specific task variables (e.g. the sensory stimulus, Fig. 4a, bottom) are related to neural activity while separating them from the impact of other factors, such as licking movements, which are captured by other model variables.

To assess how well the model predicts cortical activity, we first computed the tenfold cross-validated R^2^ (cvR^2^). For all PyN types, the model successfully captured a large fraction of single-trial variance throughout the cortex (Fig. 4b), confirming the validity of our approach. Predicted variance was higher in IT and PT mice compared to EMX mice, with over 90% explained variance in frontal cortex of PT mice. Consistent with earlier results^32,37^, movements consistently captured more variance than task variables across all PyNs (Fig. 4c). The impact of movements was also apparent when computing the unique contributions from movement or task variables by removing either set of variables from the full model, demonstrating that movements are an important driver of cortex-wide activity in all PyN types (Supp. Fig. 8a). Interestingly, unique contributions of many task variables were higher for IT, further indicating that they might be more functionally diverse than PT neurons (Supp. Fig. 8b).

**Figure 4.**
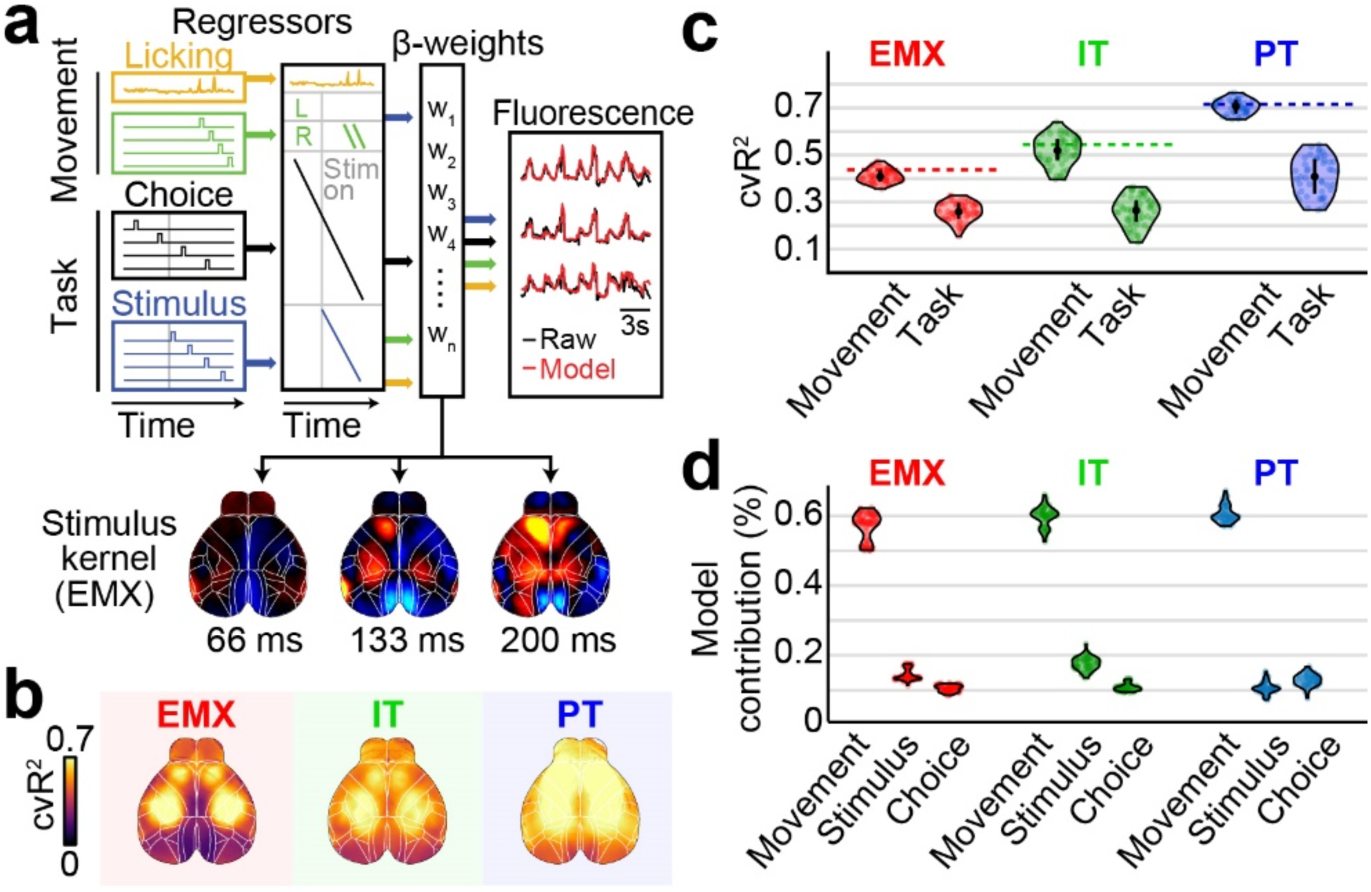
An encoding model uncovers task-specific differences across PyN types. **a)** Schematic of the encoding model. Top: Task- and movement-variables account for fluctuations in cortical activity. Bottom: Weights for each variable define a spatiotemporal event kernel, revealing cortical activity in response to a specific event (example shows right stimulus kernel in EMX mice). **b)** Average maps of cvR^2^. The model accurately predicted cortical variance for all PyNs. **c)** cvR^2^ from two models, using only movement (‘Movement’) or task variables (‘Task’). In all groups, movements were more predictive than task variables and accounted for the majority of the full models explained variance (dashed lines). Circles denote sessions. **d)** Contributions of movements, stimulus and choice to the model’s total explained variance. Although movements contributed the most, stimulus and choice also made sizable contributions.

To gain insight into the cortical dynamics related to stimulus and choice, we focused on their respective event kernels learned by the model for each PyN type. To ensure that both task variables still accounted for a sizable amount of the observed neural activity (instead of all variance being explained by movements), we first computed how much variance the stimulus and choice kernels contribute to the full model compared to the sum of all movement variables (Fig. 4d). Here, although the combined movement variables made the largest contributions (∼60% of the model’s total explained variance), both stimulus and choice also made sizable contributions to the model (10-20% explained variance in all PyN types). This demonstrates that stimulus and choice remain important for understanding cortical activity patterns and can be leveraged to selectively isolate task-related activity.

We then focused on cortical responses to the auditory stimulus. In contrast to trial averages of ΔF/F (Fig. 3c), the stimulus kernels in EMX mice uncovered clear sensory-locked and lateralized responses in auditory, parietal and frontal cortex (Fig. 4a, bottom; Fig. 5a, top). Other areas, such as somatosensory and visual cortex, were inhibited. Sensory-locked responses were also present in auditory, parietal and frontal cortices of PT and IT mice (Fig. 5a, center and bottom). However, the cortex-wide response patterns were not identical: for instance, no inhibition was apparent in PT mice. Sensory responses were particularly distinct in the parietal cortex (Fig. 5b). Responses in EMX and IT mice were more anterolateral in parietal area A while PT responses were surprisingly prominent and more posteromedial at the border between area AM and RS.

**Figure 5.**
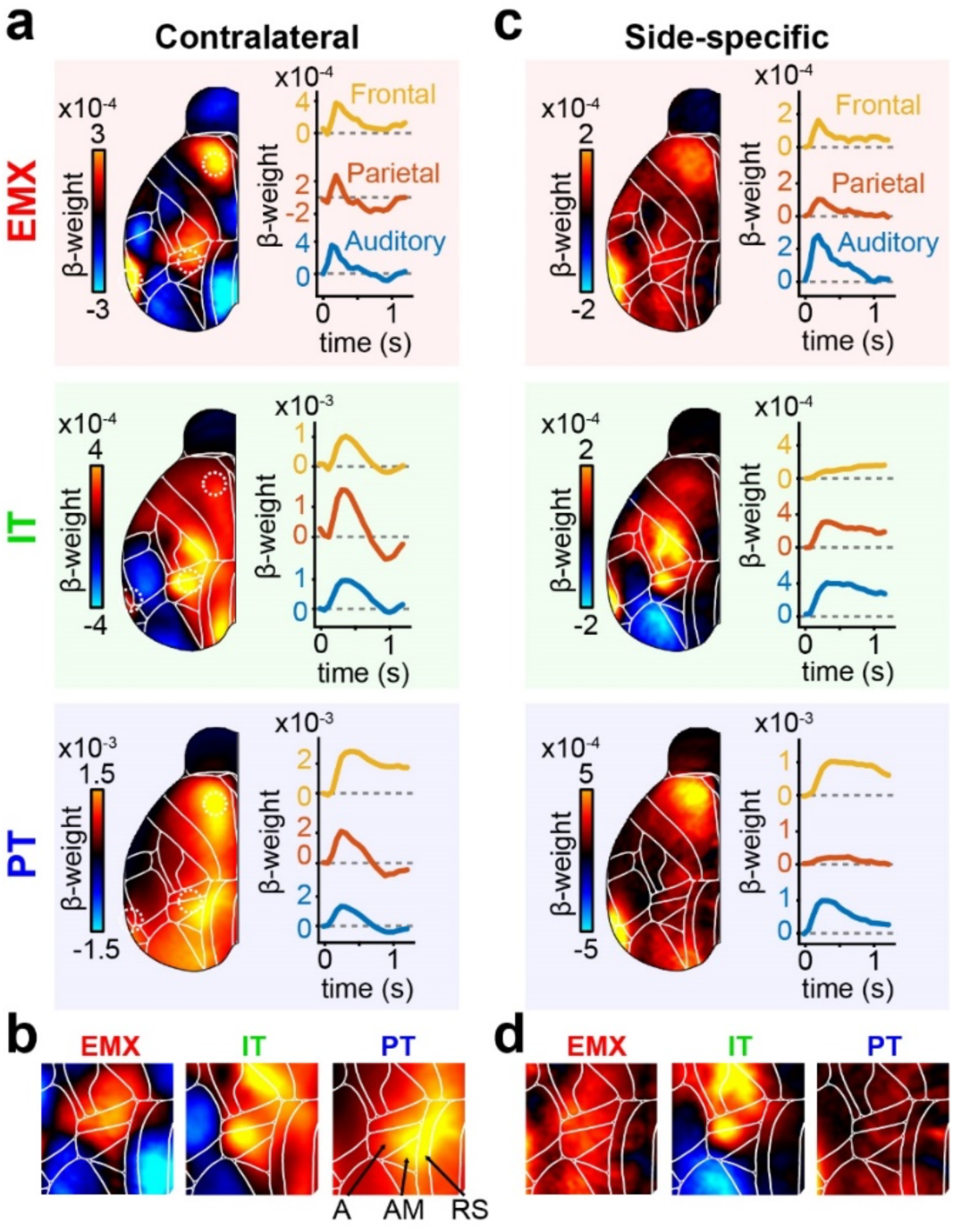
PyN-specific differences are evident in the location and specificity of cortical stimulus responses. **a)** Left: Response kernels for contralateral stimuli over all EMX (red), IT (green), and PT mice (blue), averaged between 0 and 200 ms. Right: Stimulus-evoked activity in auditory (blue), parietal (red), and frontal cortex (yellow). Dashed circles in the left stimulus maps show locations of respective cortical areas. **b)** Magnified view on parietal regions of the stimulus maps in (a). PyNs differed in the location of sensory responses. Arrows show location of parietal areas A, AM and the RS. **c)** Side-specific stimulus responses, computed as the difference between contra- and ipsilateral stimulus kernels. Conventions as in (a). **d)** Magnified view on parietal regions of side-specific maps in (c). IT neurons show clear, side-specific responses that were weaker in EMX and absent for PT neurons.

PyN types also differed in their response dynamics following the sensory stimulus. While some cortical areas, such as auditory cortex, preferentially responded to contralateral stimuli, PT neurons in parietal cortex were activated bilaterally in response to ipsi- or contralateral stimuli. To assess such side-specificity, we computed the difference between the contra- and ipsilateral stimulus kernels (Fig. 5c, colors indicate regions for which responses differed for contra-versus ipsilateral stimulus kernels). In EMX mice, we found lateralized responses in auditory, frontal, and to a lesser extent, parietal cortex (Fig. 5c,d). Lateralized IT activity was found in auditory and parietal but not frontal cortex. In contrast, PT mice showed clear side-specific responses in auditory and frontal but not in parietal cortex. Such differences in unilateral versus bilateral responses in PT and IT neurons may also reflect different functional roles, with unilateral responses encoding the spatial location of sensory information and bilateral responses signaling its importance for guiding subsequent decisions.

Having identified PyN-type dependent activity for sensory stimuli, we then examined choice-dependent activity and again observed clear differences across PyN types. In EMX mice, a number of regions showed choice-related activity, particularly in the frontal cortex, while sensory and parietal regions were only weakly modulated (Fig. 6a). We also found choice signals in somatosensory areas of the whiskers and nose that slowly increased over the course of the trial, even before the stimulus onset (Supp. Fig. 9a). These signals could indicate that mice used their whiskers to probe the location of the spouts or performed subtle, choice-predictive facial movements throughout the trial^40^. In contrast, choice-specific activity in frontal cortex strongly increased after stimulus onset and remained elevated as the decision progressed from sampling the stimulus to the subsequent delay period (Fig. 6a, yellow trace). We found equally prominent choice signals in frontal cortex of PT mice while little choice-predictive activity was seen in IT mice (Fig. 6b, Supp. Fig. 9b,c). In EMX and PT mice, positive signals for contralateral choices were concentrated in the medial part of secondary motor cortex (M2) while parts of the primary motor cortex (M1) were inhibited. This could indicate accumulation of sensory evidence and motor preparation in M2, and inhibition in parts of M1 when early lick responses must be witheld^41^.

**Figure 6.**
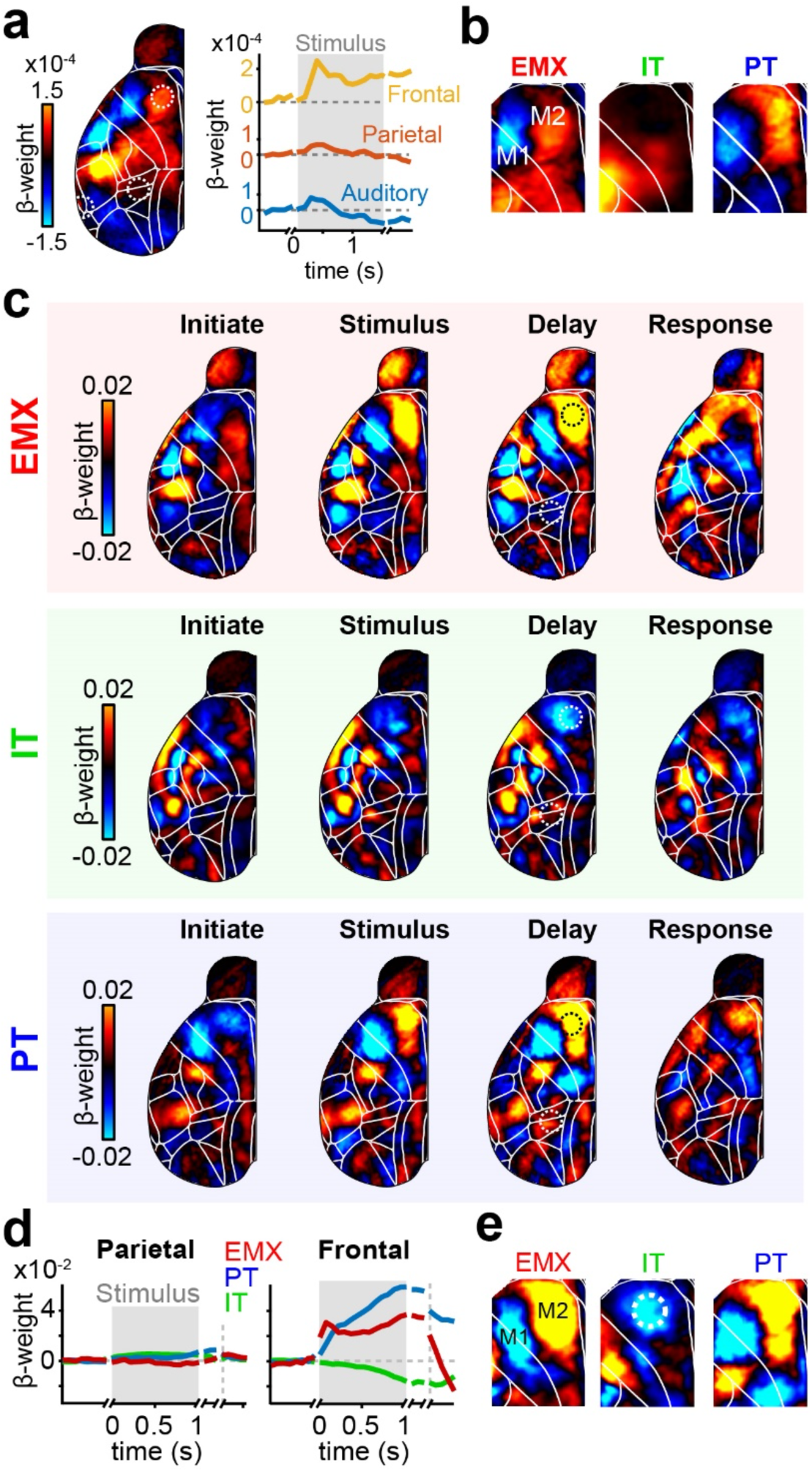
The temporal dynamics of choice-related activity differ across PyN types. **a)** Left: Averaged choice kernels for EMX mice during the delay period. Positive weights indicate increased activity for contralateral choices, negative weights indicate choice-related reduction in activity. Right: Choice-related activity in auditory (blue), parietal (red), and frontal cortex (yellow). Traces are re-aligned to the initiation, stimulus, delay and response periods, indicated by gaps in weight traces. **b)** Zoomed-in map for frontal choice kernels of EMX, IT and PT mice during the delay period. **c)** Cortical maps of contralateral choice weights for different trial episodes. Several areas in anterior cortex showed clear choice signals. **d)** Baseline-corrected decoder weights in parietal (left) and frontal cortex (right) throughout the trial. Conventions as in (a). Dashed circles in the delay maps of (c) show the parietal and frontal locations that were used to compute the traces. **e)** Zoomed-in map for frontal decoder weights of EMX, IT and PT mice during the delay period. Dashed circle shows location of ALM.

Although the choice kernels revealed differences between PyN types, choice-related activity only accounted for a small amount of the total neural variance (Fig. 4d, Supp. Fig. 8). Since the encoding model is designed to capture as much variance as possible, we hypothesized that this approach might miss subtle choice signals that are specific but low in magnitude. The linear model’s ridge penalty also enforces choice-related variance to be distributed over all correlated model variables, which might ‘diffuse’ weaker choice-related activity from the choice to other model kernels.

To selectively isolate all choice-related activity, we therefore built a decoding model, using a logistic regression choice classifier with L1 penalty (see Methods). In contrast to the encoding model, this decoder approach isolates the cortical signals that are best suited to predict the animal’s choices, regardless of their magnitude. For all PyN types, the decoder predicted the animal’s choices with high accuracy, confirming that cortical activity reliably reflects trial-by-trial choices (Supp. Fig. 10a). When analyzing the decoder weights, we found comparable patterns to the encoding model’s choice kernels but with much clearer separation of cortical areas (compare top row ‘Delay’ in Fig. 6c to Fig. 6a, left). Here, positive decoder weights denote areas that are most predictive for contralateral choices but, importantly, this does not suggest that these areas are necessarily the most active. For all PyN types, we found significant choice signals in multiple areas of the anterior cortex that evolved during decisions (Fig. 6c, Supp. Figs. 10, 11). In EMX and PT mice (top and bottom rows), large parts of M2 were again highly choice predictive, including the anterior lateral motor cortex (ALM) and the medial motor cortex (MM)^21^. M2 choice weights strongly increased immediately after stimulus onset and remained elevated during the subsequent delay period (Fig. 6c,d). Cortical choice signals also persisted after removing movement-related activity from the data, suggesting that they are not explained by choice-predictive animal movements but instead reflect the formation of sensory-driven decisions in frontal cortex (Supp. Fig. 12).

Surprisingly, we also found a mild ipsilateral choice-preference for M2 in IT mice, despite strong bilateral activation of frontal cortex during the delay period (Fig. 3c; Supp. Fig. 7). This ipsilateral choice signal evolved more slowly during the stimulus and delay period (Fig. 6c,d) and was more spatially restricted to ALM (Fig. 6e). In line with our previous results, no choice signals were seen in parietal cortex of any PyN type (Fig. 6d, left), suggesting that parietal cortex is mostly involved in sensory processing instead of choice formation or motor execution^42,43^.

Compared to the encoding model, the decoding model recovered more finely structured choice maps, especially in the frontal regions (compare Fig. 6e to 6b), revealing strong contralateral choice signals in PT and ipsilateral choice signals in IT mice. One possible cause of this unexpected inversion may be different choice-selectivity of specific IT-subtypes: intracortical versus corticostriatal (CStr) projection neurons. Earlier work suggested an even distribution of ipsi- and contralateral choice selectivity in frontal intracortical projection neurons^20,21^, and we thus hypothesized that our observed IT choice selectivity is due to the activity of CStr neurons. To address this directly, we used an intersectional approach that exclusively labels large populations of CStr neurons by performing multiple bilateral striatal injections of a retrograde virus expressing Cre (CAV2-Cre) in GCaMP6s reporter mice (Fig. 7a), inducing widespread expression in CStr neurons (Fig. 7b). As expected, GCaMP6 expression in CStr neurons was largely confined to layer 5 with sparse expression in deeper layers.

**Figure 7.**
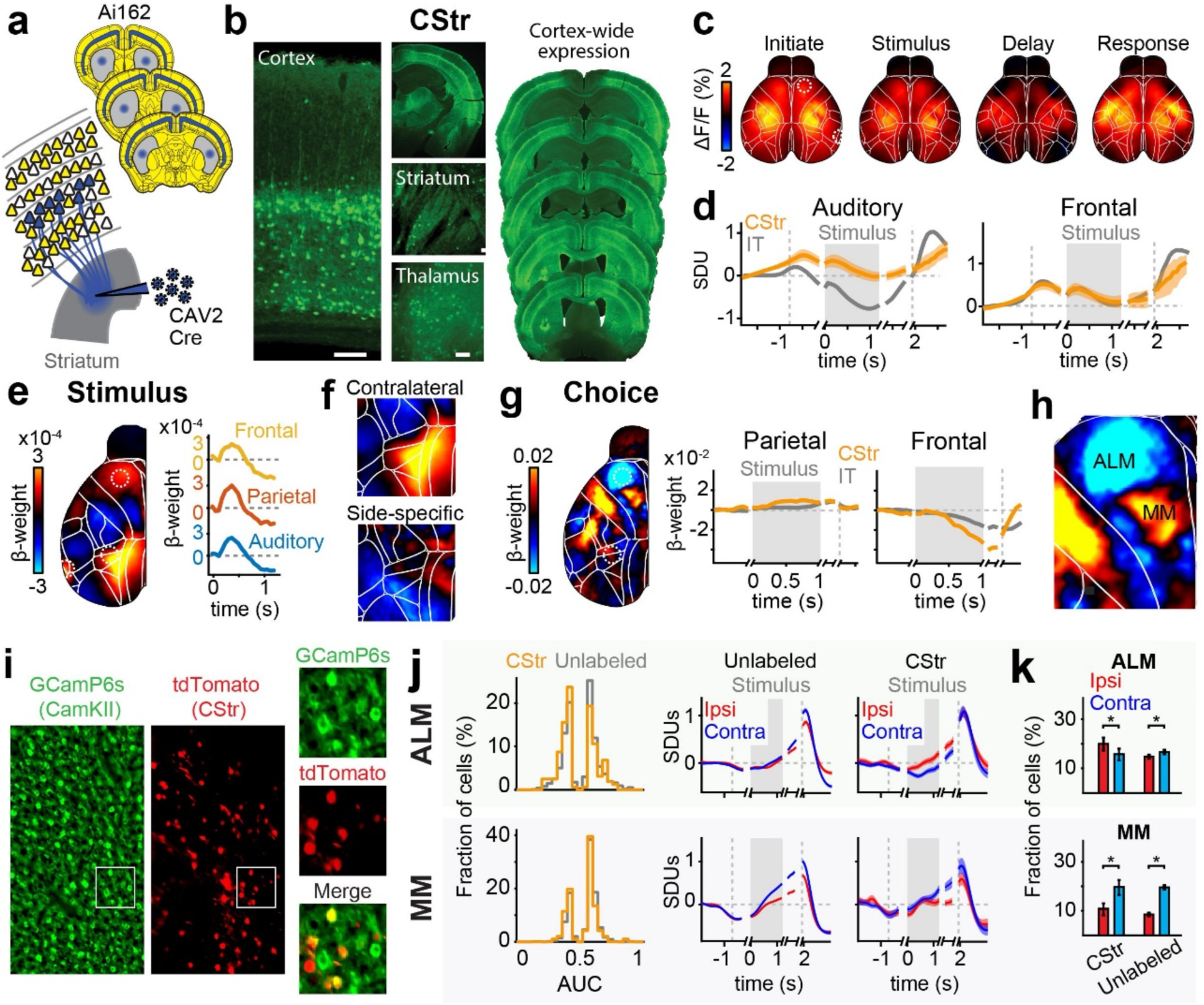
An intersectional approach to measure cortex-wide activity of CStr neurons. **a)** Schematic of the retrograde labeling approach. Multiple bilateral injections of retrograde CAV2-Cre virus in reporter mice robustly induced expression of GCaMP6s in CStr neurons. **b)** Left: GCaMP6s-expression was robust throughout the dorsal cortex. Scale bars are 100 µm. Right: Brain slices show robust cortex-wide expression. **c)** Cortical dynamics of CStr neurons in the auditory discrimination task. Shown are trial-averages over all correct, leftward trials in different trial episodes. No clear lateralization was observed in trial averages. **d)** Averaged activity in auditory (left) and frontal cortex (right) for CStr (orange) and IT (gray) mice. Dashed lines show times of initiation and response periods, gray areas the stimulus period. Traces show standard deviation units (SDU). **e)** Left: Contralateral stimulus kernel, averaged over all CStr mice, between 0 and 200 ms after stimulus onset. Right: Colored traces show changes in sensory (blue), parietal (red) and frontal cortex (yellow). Locations for each area are indicated by dashed circles in the weight map. **f)** Top: Weight map from (e), zoomed in on parietal cortex. Bottom: Difference of contra-versus ipsilateral stimulus kernels. **g)** Left: Cortical weight map from choice decoder during the delay period, averaged over all CStr mice. Right: Baseline-corrected decoder weights in parietal (left) and frontal cortex (right) for CStr (orange) and IT mice (gray). Traces are re-aligned to the initiation, stimulus, delay and response periods, indicated by gaps in weight traces. **h)** Weight map from (g), zoomed in on frontal cortex. **i)** Example field-of-view from two-photon imaging. Left panel: GCaMP6s-expression in all PyNs (green). Center panel: retrogradely-labeled CStr neurons, expressing tdTomato (red). Right panels: zoomed-in maps for both channels and a merged image. **j)** Left: Overview of significantly choice-tuned neurons in ALM (top) and MM (bottom), imaged between 200-400 µm. Orange line shows CStr neurons, gray lines are unlabeled PyNs. AUC values below 0.5 indicate stronger responses for ipsilateral choices. Right: Trial-averaged activity for all choice-selective neurons, separated for ipsi- (red) versus contralateral choices (blue). CStr neurons in ALM (top right) show higher activity for ipsilateral choices. **k)** Fraction of cells responding selectively for ipsi-versus contralateral choices. Top: A higher fraction of CStr neurons in ALM are ipsi-selective (CStr_Ipsi_: 20.4%, CStr_Contra_: 15.5%, binomial test: *p* = 0.0018) while unlabeled neurons are more contra-selective (Unlabeled_Ipsi_: 14.3%, Unlabeled_Contra_: 17.2%, binomial test: *p* = 3.5×10^−10^). Bottom: The majority of CStr as well as unlabeled neurons in MM are selective for contralateral choices (CStr_Ipsi_: 10.2%, CStr_Contra_: 19.1%, binomial test: *p* = 2.7×10^−8^; Unlabeled_Ipsi_: 9.3%, Unlabeled_Contra_: 19.6%, binomial test: *p* < 1×10^−10^).

We then used widefield calcium imaging to selectively measure activity from CStr neurons. As with PT and IT mice, we observed robust fluorescence signals throughout the dorsal cortex (Supp. Movie 5) and identified visual areas using retinotopic mapping (Supp. Fig. 13a). sNMF showed that the dimensionality of CStr mice was intermediate between PT and IT activity, and that the spatial components did not strongly overlap with other PyN types (Supp. Fig. 13b,c). This clear difference between IT and CStr mice strongly suggests that imaging signals from IT mice are not solely dominated by IT-positive CStr neurons but represent a mixture of intracortical- and corticostriatal-projecting IT neurons with distinct activity patterns.

We then trained CStr mice in the auditory discrimination task. Trial-averaged dynamics partially resembled IT mice, for example in frontal cortex, but also showed clear differences, such as a lack of pre-stimulus suppression in sensory cortex (Fig. 7c,d; orange versus gray traces). Differences between CStr and IT mice were also visible in the stimulus kernels from the encoding model. In CStr mice, stimulus-related activity in parietal cortex was stronger than in sensory and frontal cortex but the activated parietal regions were more medial than in IT mice (Fig. 7e). Interestingly, these stimulus-driven parietal regions (Fig. 7f, top) closely resembled cortical areas that form anatomical and functional connections to the dorsomedial striatum^44,45^. As with PT neurons (Fig. 5b,d), parietal CStr responses equally responded to contra- and ipsilateral stimulation (Fig. 7f, bottom).

To determine whether CStr activity contributed to ipsilateral-preferring IT choice signals, we repeated the choice decoder analysis. The decoder predicted animals’ choices with equally high accuracy as for PT and IT mice (Supp. Fig. 13d). We then extracted choice weights for each task episode. Here, CStr activity was overall similar to that of IT mice, with an even stronger ipsilateral-choice preference in frontal cortex that started after stimulus onset and lasted throughout the delay and response periods (Fig. 7g, right). As in IT mice, this inversion from contra- to ipsilateral choice preference was prominent in area ALM but did not extend to MM, strongly suggesting that ipsilateral choice preference is driven by IT-CStr neurons.

To confirm these results on a single-cell level, we then used two-photon calcium imaging in Camk2α-tTA;G6s2 mice and labeled CStr neurons through striatal injections of a retrograde adeno-associated virus (AAVrg-tdTomato) (Fig. 7i). This allowed us to compare the activity of all PyNs in frontal cortex with tdTomato-labeled CStr neurons in the same animal. Comparing the choice-tuning of CStr and unlabeled PyNs revealed a specific difference in ipsi-versus contralateral choice preference in ALM (Fig. 7j). Here, significantly more choice-selective CStr neurons preferred ipsilateral choices, whereas unlabeled PyNs were mildly contra-selective (Fig. 7k). In agreement with our widefield results, these differences were only seen in ALM but not MM. Interestingly, this ipsilateral choice preference was also restricted to superficial IT-CStr neurons at a cortical depth between 200-400 µm. Infragranular CStr neurons (400-600 µm), which are also often PT cells^17^, showed strong contralateral choice tuning that was similar to unlabeled cells (Supp. Fig. 14a). Lastly, we also tested if the neuropil exhibits distinct choice signals that may have masked somatic activity in our PyN-specific widefield measures (Supp. Fig. 14b). In all regions and depths, neuropil choice tuning was largely similar to somatic activity of unlabeled neurons, confirming that widefield measures indeed represented local somatic activity. Here, IT-specific widefield signals matched the mixed choice-selectivity of superficial layers, while PT-specific imaging was well-aligned with the clear contralateral choice tuning in deeper cortical layers.

The observed differences between PyN types suggest that each type may drive distinct aspects of decision-making. To causally test the functional role of different PyN types, we performed PyN-specific optogenetic inactivation in different cortical areas. We induced Cre-dependent expression of the inhibitory opsin stGtACR2^46^ by injecting an AAV in the parietal and frontal cortex of Fezf2-, PlexinD1-, or EMX-Cre mice (Fig. 8a, b, Supp. Table 2). To express stGtACR2 in CStr neurons, we used a combined viral receptor complementation and intersectional approach to maximize the efficiency of retrograde expression^47^. First, we simultaneously expressed the coxsackievirus and adenovirus receptor (CAR) and transduced Cre-dependent stGtACR2 in cortical neurons. Two weeks later, we performed retrograde-mediated stGtACR2 expression through striatal CAV2-Cre injections. CAR mediates efficient axonal uptake of CAV2 and prevents potential viral tropism in subsets of CStr neurons. The coordinates of auditory, parietal, and frontal injections and subsequent inactivation were determined from our stimulus and choice analyses (Fig. 5a, 6a; dashed circles). To test whether optogenetic effects are area-specific, we also targeted the primary visual cortex (V1) in a subset of EMX mice as controls. To simplify the behavioral task during the optogenetic inactivation experiments, we only presented unilateral stimuli in each trial and kept the stimulus duration and delay periods constant (1 and 0.5 seconds, respectively).

**Figure 8.**
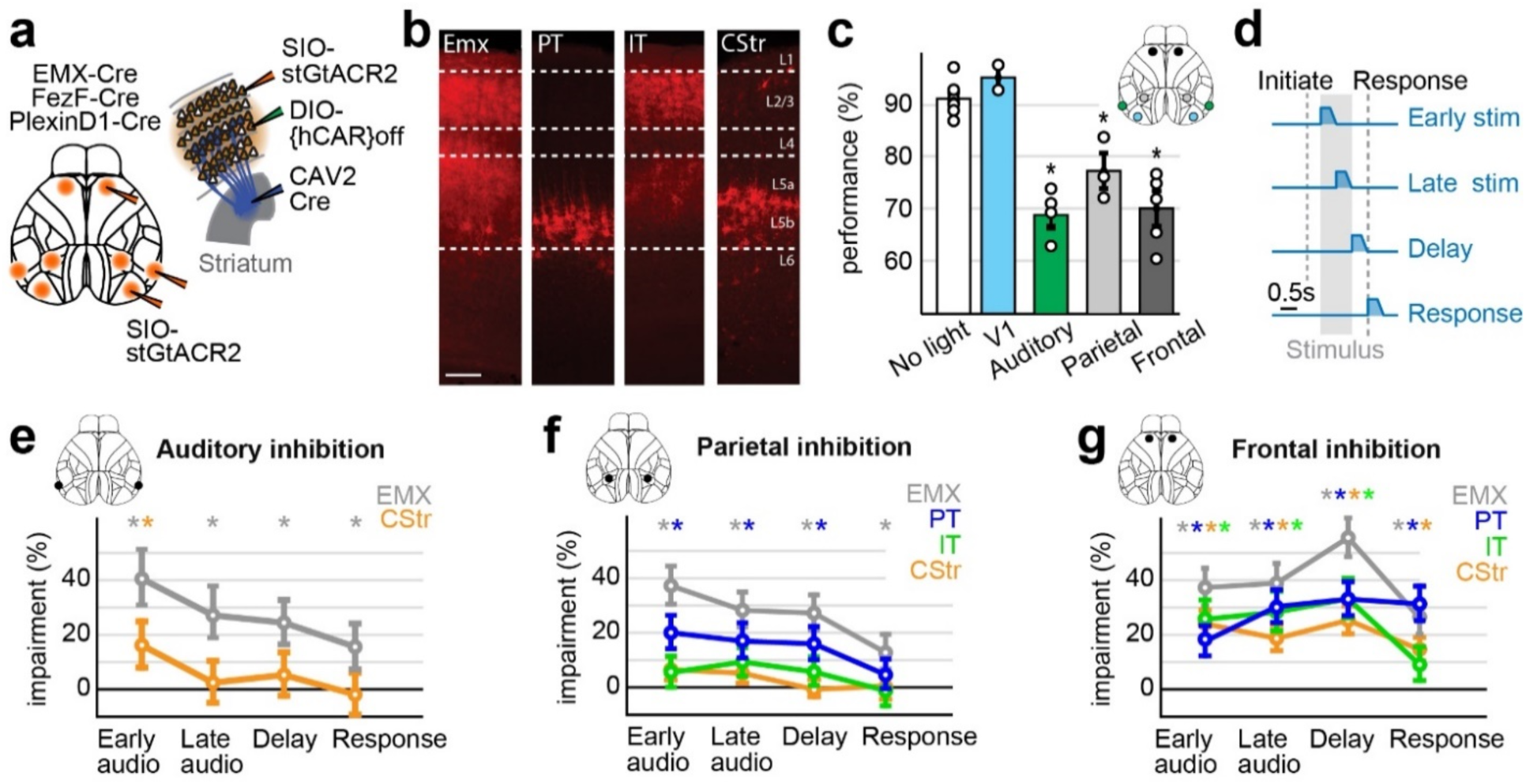
Temporally restricted, PyN-specific inactivation of parietal and frontal cortex disrupts decisions. **a)** Left: Schematic of injection scheme to induce stGtACR2 expression in EMX, IT, or PT neurons. Injections were performed in auditory, parietal and frontal cortex of transgenic mice to target different PyN types. V1 injections were performed in a subset of control EMX mice. Right: Intersectional viral approach for targeting CStr neurons. AAV-DJ-hSYN-DIO-hCAR{off} induced expression of CAR in cortical neurons to promote uptake of CAV2-Cre virus from the striatum. A second cortical injection induced Cre-dependent stGtACR2 expression in all CStr neurons. **b)** Layer-specific expression of stGtACR2-FusionRed in EMX, PT, IT and CStr neurons. **c)** Behavioral performance (% correct) of EMX mice during inactivation of V1, auditory, parietal or frontal cortex. Inhibition of auditory, parietal and frontal cortex, but not V1, significantly reduced task performance. Circles indicate individual mice, error bars show the s.e.m. **d)** Schematic of optogenetic inactivation paradigm. 0.5-s long optogenetic inhibition was performed during the first or last part of the stimulus period, the subsequent delay or the response period. Light power ramped down after 0.3 seconds. **e)** Behavioral impairment (% change from control performance) with inhibition of EMX or CStr neurons in auditory cortex. Circles show mean impairments, error bars 95% confidence intervals. **f)** Behavioral impairment with inhibition of different PyN types in parietal cortex. Conventions as in e). **g)** Behavioral impairment with inhibition of different PyN types in frontal cortex. Conventions as in f). Stars in all panels indicate significance (bonferroni-corrected p < 0.01, binomial test).

In agreement with our imaging results, bilateral optogenetic inactivation of auditory, parietal or frontal cortex of EMX mice strongly impaired auditory decision accuracy (Fig. 8c). Performance was unaffected by V1 inactivation, confirming that these areas are selectively required for auditory decisions. We then illuminated each area for 0.5 seconds during four different task episodes: Early and late stimulus (the first and last 0.5 seconds of the stimulus period), delay and response (Fig. 8d). In agreement with our observation that auditory and parietal cortex reflects stimulus driven activity (Fig. 5a, red and blue traces), inactivation of either area strongly reduced animal’s performance, particularly during the stimulus period (Fig. 8e, f). Behavioral impairments (computed as the normalized difference between performance in non-optogenetic trials and chance) were weaker during the subsequent trial periods, indicating that these areas are most important for early processing of auditory stimuli. As expected, impairments were strongest in EMX mice where all PyNs were affected (gray trace).

In agreement with earlier work^48^, inhibiting CStr neurons in A1 also affected auditory decisions (Fig. 8e, orange trace). However, the effects were more transient and weaker compared to inactivation of EMX neurons, suggesting that CStr neurons were not exclusively required for accurate task performance. This was even more evident in parietal cortex where impairments were surprisingly weak when inhibiting either IT or CStr neurons (Fig. 8f, green/orange traces) while inactivating PT neurons resulted in a more robust impairment (blue trace). This suggests that the subcortical PT projection pathways in the parietal cortex have a larger causal impact on sensory processing than intracortical IT or CStr projections, pointing to a role for PT neurons that extends beyond preparation and execution of movements.

Frontal inactivation resulted in a particularly strong impairment, especially for nonspecific PyN inactivation (EMX mice, Fig. 8g, gray trace), consistent with a role for frontal cortex in sensory integration and working memory^20,43,49^. IT and CStr inactivation resulted in comparable behavioral impairments during the stimulus and delay periods (Fig. 8g, green and orange traces). Choice impairments in IT mice are therefore not solely due to the disruption of intracortical processing^20^ but also involve alterations of CStr neurons. Inactivating PT neurons equally impaired animal performance during the stimulus and delay period but showed stronger effects during the final response period. Impairments in the response period were similar for EMX and PT mice, suggesting that PT neurons are particularly involved in licking responses. These results show that multiple PyN types in frontal cortex are involved in the formation and maintenance of choices, despite clear differences in their respective choice tuning. Lastly, we also analyzed licking patterns to test if optogenetic inhibition broadly disrupted animal movements. Frontal inactivation in the delay period had a mild effect on response latency but did not affect response probability or licking patterns, arguing against a strong motor impairment (Supp. Fig. 15). This shows that PyN-specific inhibition selectively reduced the animals’ response accuracy rather than broadly disrupting their ability for movement initiation and execution.

## Discussion

We measured and manipulated PyN subtypes to determine whether they play distinct roles in decision-making. Cortex-wide activity patterns were PyN subtype-specific, each reflecting distinct neural dynamics at multiple spatial scales. Functional specificity across PyN subtypes was also evident during auditory decision-making: activity of each PyN subtype exhibited unique cortical localization and spatial specificity associated with stimulus and choice. These response patterns were not seen when imaging from PyNs nonspecifically. Optogenetic inactivation confirmed that PyN types in parietal and frontal cortex have distinct functional roles, highlighting the importance of subcortical projection pathways for sensory processing and choice formation. Taken together, our results suggest that different PyN types form parallel subnetworks throughout the cortex, are functionally-distinct, and perform separate roles during auditory decision-making.

Dimensionality reduction of cortical dynamics^33,50,51^ revealed that nearly all spatial components were also PyN-subtype-specific. Differences between PyNs are therefore not only present within parcellated cortical regions^18–20^ but also evident in large-scale activity patterns. This has important implications for studies of cortex-wide neural dynamics, which are often based on indirect measures of neural activity^52,53^ or pooled activity from all PyNs^22,25,54,55^. Earlier work revealed intricate circuit motifs and functional modules that span the entire cortex^50,54–57^ and largely follow intracortical connectivity patterns^58,59^. Our results point to the existence of additional circuit motifs when specific PyN population are isolated, especially subcortical projection types, such as PT or CStr neurons. Furthermore, most PyN-specific LocaNMF components consisted of spatially precise subregions that were smaller than Allen CCF areas. Detailed analysis of PyN-specific activity might therefore reveal more detailed cortical structures than could be observed with nonspecific measures. For future studies, a particularly promising approach to achieve this goal would be to combine large-scale measures of multiple PyN types with multi-color widefield imaging^24,60^ and directly observe interactions between PyN-specific cortical dynamics within the same animal.

Sensory responses were pronounced in auditory, parietal and frontal cortices^42,43,61,62^ and revealed unique response patterns for each PyN type. This functional segregation is in line with recent results from primary sensory areas, such as somatosensory^18,63^ and visual cortex^19^, and strongly suggests that different PyN types play separate roles during sensory processing. Moreover, the clear differences in magnitude, localization, and lateralization of sensory responses in parietal and frontal cortex demonstrate that the functional specialization of different PyN subtypes is a general feature of cortical circuit function.

We selectively inactivated different PyN types across cortical areas to directly test their involvement in the behavioral task. In agreement with earlier work, inhibiting CStr neurons in auditory cortex impaired task performance^48^, suggesting that projections from cortex to striatum are important for accurate auditory decisions. Moreover, inactivation of parietal cortex also caused strong behavioral impairments, especially during sensory stimulation. Surprisingly, inactivation of CStr or IT neurons caused only weak behavioral effects. This shows that the importance of CStr projections does not generalize from auditory to parietal cortex and also argues against models in which task-relevant sensory information is directly transmitted from parietal to frontal cortex to promote decision formation^42,62,64^. Instead, inactivation of parietal PT neurons during the stimulus presentation strongly disrupted eventual decisions, highlighting the importance of subcortical projection pathways for auditory performance. A likely subcortical target is the SC, which receives inputs from PT neurons^18,26^ and has been implicated with somatosensory^18^ and visual^65,66^ perception, and decision-making^39,67,68^.

The strong behavioral effects with inactivation of parietal cortex are at odds with earlier studies in rats that showed very little or no impact of parietal inactivation in auditory task performance^69,70^. Conversely, other work in head-fixed mice had found robust behavioral effects in visual^43,71–73^ and auditory^62^ task performance. While a potential reason for these contradictory results could be species differences between rats and mice, the precise location of parietal inactivation most likely plays an important role. Different sensory modalities are processed along a mediolateral gradient in parietal cortex, emphasizing the need to precisely target specific parietal areas to obtain a modality-specific behavioral effect^62^. We also found that parietal responses of PT and CStr neurons were located more medially than IT neurons, further emphasizing the need for precise localization. Another important factor is the specific behavioral task. Our task design required accumulation of sensory evidence and working memory which engage a wider range of cortical regions^72^ and might therefore explain the involvement of parietal cortex for accurate auditory decisions.

The accumulation and memory requirements of the decisions we studied might also explain why we found clear cortical choice signals, especially in frontal cortex, whereas recent cortex-wide studies reported little choice-selectivity^39,74^. The lack of side-specific choice tuning in IT population data is consistent with earlier work, showing that intracortical projection neurons in frontal cortex include an equal amount of contra- and ipsilateral choice-preferring cells^20,21^. In contrast to other PyNs, CStr neurons were more selective for ipsilateral choices. Using two-photon imaging, we confirmed that this ipsilateral choice preference was also present in individual CStr neurons but not in unlabeled IT neurons. This confirms that our PyN-specific widefield results indeed reflect selective somatic activity and not just superficial neuropil signals, in line with earlier findings that widefield signals are strongly correlated to spiking of cortical neurons^44^. Ipsilateral choice signals in CStr neurons were restricted to superficial layers of area ALM, which is mostly implicated in movement generation^13,49^. A recent study showed that CStr projections from the anterior cingulate cortex can inhibit striatal activity and motor behavior^75^. A potential reason for the reduced activity of CStr neurons during the response period could therefore be to disinhibit striatal circuits and release a targeted licking response.

In agreement with our imaging results, frontal inactivation strongly impaired animal behavior during the stimulus and delay period, suggesting an important role for the translation of sensory inputs into behavior^21,74,76–78^. Impairments were largely similar for all frontal PyN types, which appear to be equally required for choice formation and retention. Frontal PyNs may thus be more reliant on each other to maintain accurate function than in sensory areas^18,19,48^. As the only exception, PT neurons were more important during the response period, consistent with a specific role of brainstem-projecting PT neurons for motor execution^13^.

Our work offers a new perspective on cortex-wide dynamics by viewing them through the lens of different PyN types and strongly supports the view that cortical circuits perform parallel computations, even within the same cortical layer^13,18,19,79^. Future work to reveal the circuit logic of cortex-wide activity patterns in different behavioral contexts should therefore include PyN type information to resolve the functional heterogeneity that is often encountered when studying cortical decision circuits. A powerful tool to achieve this goal are novel mouse lines, such as our temporally inducible knock-in lines that allowed us to reliably target PT and IT neurons with high specificity. These new mouse lines also overcome several known problems of earlier transgenic approaches, unstable expression patterns across cortical areas or mixtures of cell types due to interactions with surrounding genetic elements^26,80^. Future studies could explore PyN type function at even higher granularity by using more specific mouse lines, such as the Cux1-line to selectively isolates intracortical projection neurons^26^. Moreover, combining genetic mouse lines with retrograde labeling will enable the targeting of specific PyN subtypes, such as projection-specific PT neurons^12,13^, that might serve a large array of functions from sensory processing, to working memory and motor function^13,18^.

## Acknowledgements

We thank Z. Josh Huang for providing the Fezf2-2A-CreER and PlexinD1-2A-CreER mouse line and for countless insightful conversations; Shreya Saxena and Joao Couto for technical advice and scientific discussions; Anup Khanal for assistance with animal perfusions; Sandra Brill, Maria Perez and Alice Despatin for animal training; Adam Kepecs for providing the viral vector AAV-DJ-hSYN-DIO-hCAR{off}; Katherine S. Matho for invaluable help with Fezf2 and Plexin cell density quantification. This work was supported in part by NIH grants R01EY022979 and W911NF-16-1-0368 as part of the collaboration between the US DOD, the UK MOD and the UK Engineering and Physical Research Council under the Multidisciplinary University Research Initiative to A.K.C. S.M. was funded by the Deutsche Forschungsgemeinschaft (DFG, German Research Foundation) - 368482240/GRK2416 and the Helmholtz association (VH-NG-1611). X.R.S. was supported by the NRSA F32 MH120888.

## Methods

### Mouse lines

All surgical and behavioral procedures conformed to the guidelines established by the National Institutes of Health and were approved by the Institutional Animal Care and Use Committee of Cold Spring Harbor Laboratory. Experiments were conducted with male mice between the ages of 8 to 25 weeks. No statistical methods were used to pre-determine sample sizes but our sample sizes are similar to those reported in previous publications^22,25^. All mouse strains were acquired from the Jackson Laboratory, the Allen Brain Institute, or generated at Cold Spring Harbor Laboratory. Transgenic strains crossed to generate double- and triple-transgenic mice used for imaging: Emx-Cre (JAX 005628), LSL-tTA (JAX 008600), Ai93D (JAX 024103), Ai162 (JAX 031562), TRE-GCaMP6s (G6s2, JAX 024742) and H2B-eGFP (JAX 006069). EMX mice, used for calcium imaging, were bred as Ai93D;Emx-Cre;LSL-tTA. To avoid potential aberrant cortical activity patterns, EMX mice were on a doxycycline-containing diet (DOX), preventing GCaMP6s expression until they were 6 weeks or older^22,25^.

For widefield imaging of PT and IT neurons, inducible knock-in drivers Fezf2-2A-CreER and PlexinD1-2A-CreER (Supp. Table 2), respectively, were crossed with Ai162 reporter mice to drive cortex-wide GCaMP6s expression. Cre activity was induced through two doses of intra-peritoneal injections of tamoxifen (200 mg/kg; 20 mg/ml solution dissolved in corn oil) at P28 and P32. Histologic characterization revealed pyramidal neuron expression patterns consistent with prior reports^26^. For widefield imaging of corticostriatal neurons, we first crossed Ai162 with G6s2 to create a new double-transgenic reporter strain Ai162;G6s2 with two hemizygous copies of GCaMP6s under tetO control. Since LSL-tTA is incorporated in-tandem to the reporter gene in the Ai162 strain^29^, this hybrid reporter line permits Cre-dependent expression of GCaMP6s at higher levels than Ai162 hemizygotes while avoiding potential leaky reporter gene expression that is sometimes found in Ai162 homozygotes. Next, to achieve widespread GCaMP6s expression in corticostriatal neurons, we performed striatal injections of retrograde virus (CAV2-Cre) in the hybrid Ai162;G6s2 reporter line (see section: viral injections).

For two-photon imaging experiments, GCaMP6s expression in pyramidal neurons were generated using the hybrid strain Camk2α-tTA;G6s2.

### General surgical procedures

All surgeries were performed under 1-2% isoflurane in oxygen anesthesia. After induction of anesthesia, 1.2 mg/kg of meloxicam was injected subcutaneously and sterile lidocaine ointment was applied topically to the skin incision site. After making a midline cranial incision, the skin was retracted laterally and fixed in position with tissue adhesive (Vetbond, 3M). We then built an outer wall using dental cement (C&B Metabond, Parkell; Ortho-Jet, Lang Dental) along the lateral edge of the dorsal cranium (frontal and parietal bones) to maximize the area of exposed skull. A custom titanium skull post was then attached to the dental cement. For skull clearing, debris and periosteum were thoroughly cleaned from the skull followed by the application of a thin layer of cyanoacrylate (Zap-A-Gap CA+, Pacer technology).

To prepare mice for two-photon imaging, a circular craniotomy (ø = 3 mm) centered over the right frontal cortex (1.75 mm lateral to midline and 1.75 mm rostral to bregma), was made using a biopsy punch. A circular coverslip (ø = 3 mm) was then lowered to the surface of the brain and Vetbond and Metabond were used to seal the window to the skull. Lastly, a titanium skull post was implanted as described above.

### Viral injections

After induction with isoflurane anesthesia, animals were placed in a stereotaxic frame (David Kopf Instruments). The skull was leveled along the rostral-caudal and medio-lateral axis, allowing for accurate and reproducible targeting. All injections were made using a programmable nanoliter injector (Nanoject III, Drummond Scientific, PA). For *widefield imaging* of CStr mice, widespread corticostriatal GCaMP6s expression was generated in Ai162;G6s2 reporter mice by performing bilateral stereotaxic injections of CAV-2-Cre (at 3-4 weeks of age) into the dorsal striatum at three targets per hemisphere spanning the rostro-caudal axis. The target coordinates (relative to bregma and dura, in mm) are (1) RC +0.75, ML ±1.8, DV 3.0; (2) RC 0, ML ±2.2, DV 3.1; (3) RC −0.75, ML ±2.9, DV 3.1. For each striatal target, a burr hole was created using a small dental burr followed by injection of 1.8×109 purified particles (pp) of CAV-2-Cre using pipettes with long taper tips pulled from borosilicate capillaries (3.5” Drummond # 3-000-203-G/X, Drummond Scientific, PA). For *two-photon imaging* experiments, CStr neurons were labeled through striatal injections of AAV-2-retro-CAG-tdTomato (using the same approach and coordinates as described above) in Camk2α-tTA;G6s2 mice.

For cell type-specific *optogenetic silencing* experiments, we performed bilateral injections in frontal, parietal, and auditory cortex (coordinates relative to bregma, in mm: frontal cortex: RC +2.5, ML ±1.5; parietal cortices: RC −1.7, ML ±2.5; auditory cortex: RC −2.5, ML ±4.6) to induce expression of Cre-dependent stGtACR2 (AAV1-hSyn-SIO-stGtACR2-FusionRed, Upenn Vector Core, Supp. Table 2). Cortical injections were performed in P42 to P56 Fezf2-2A-CreER (PT), PlexinD1-2A-CreER (IT), and EMX-Cre (nonspecific PyNs) reporter mice. In mice with ligand-dependent Cre recombinase activity, intraperitoneal tamoxifen was administered one week after viral injections. Injections were made at two depths (300 and 600 µm) per cortical target. In two EMX-Cre mice, bilateral injections were performed in the frontal and visual cortex (RC −4, ML ±2.5). To express stGtACR2 in CStr neurons, viral injections were performed in C57BL/6J mice in two stages. In stage one, we utilized a viral receptor complementation strategy^47^ by injecting both AAV-DJ-hSYN-DIO-hCAR{off} and AAV1-SIO-hSyn1-stGtACR2-FusionRed in different cortical locations (targets as described above) in P21-P28 mice. In stage two, we performed bilateral striatal CAV-2-Cre injections 6 weeks after the initial cortical injections (see above). hCAR is expressed in all transfected neurons in a Cre-OFF manner, meaning that Cre expression stops expression of hCAR while inducing expression of stGtACR2.

### Optical fiber implantation

For optogenetic silencing, we expressed the soma-targeted anion-conducting channelrhodopsin stGtACR2 in target neuronal populations^46^. Optical fibers (NA = 0.36, ø = 0.4mm, FT400UMT, Thorlabs) were glued into metal or ceramic ferrules (ø = 1.25 mm, Thorlabs) and secured above the cortex following viral injections at each target. Ferrule-enclosed optical fiber implantations immediately followed cortical AAV injections in Fezf2, PlexinD1, and Emx mice and striatal CAV-2-Cre injections in CStr mice. One polished end of the optical fiber was positioned extradural to the site of cortical injections and interfaced with thinned skull using cyanoacrylate. Next, the fiber was fixed to the skull using light-cured glass ionomer (Vitrebond, 3M). Additional layers of dental cement and dental acrylic (Lang Dental Jet Repair Acrylic; Part#1223MEH) were applied around the fiber implant and the skull to reinforce for durability and long-term stability. After all layers were cured, a final outer coating of cyanoacrylate and nail polish were applied.

### Behavioral training

The behavioral setup was controlled with a microcontroller-based (Arduino Due) finite state machine (Bpod r0.5, Sanworks) using custom Matlab code (2015b, Mathworks) running on a Linux PC. Servo motors (Turnigy TGY-306G-HV) and touch sensors were controlled by microcontrollers (Teensy 3.2, PJRC) running custom code. Fifty-four mice were trained on a delayed, spatial discrimination task. Mice initiated trials by placing their forepaws on at least one of the two handles, which were mounted on servo motors that rotated out of reach during the inter-trial period. Upon trial initiation, animals placed their forepaws on the handles and, after a variable duration of 0.25-0.75 seconds of continuous contact, the auditory stimulus was presented. Auditory stimuli consisted of a sequence of Poisson-distributed, 3-ms long auditory click sounds^36^, presented from either a left and/or right speaker for a variable duration between 1 and 1.5 seconds. The stimulus period was followed by a variable delay of up to 1 second, then the servo motors moved two lick spouts into close proximity of the animal’s mouth. If the animal licked twice on the side where more clicks were presented, a drop of water reward was dispensed. The amount of water rewarded per trial (typically 1.5 to 3 µl) was constant within a single session but was sometimes adjusted daily based on the animal’s body weight. After one spout has been licked twice, the contralateral spout moved out of reach to force the animal to commit to its decision.

All trained mice were housed in groups of two or more under reverse light cycle (12-hour dark and 12-hour light) and trained during their active dark cycle. Animals were trained over the course of approximately 30-60 days. After 2-3 days of restricted water access, animals began habituation to head fixation and received water from spouts in the behavior chamber. During these sessions, unilateral auditory stimuli were presented followed by a droplet of water dispensed freely from the ipsilateral water spout. After several habituation sessions, animals were then required to touch the handles to trigger stimulus presentation. Once mice could reliably reach for the handles, the required touch duration was progressively increased up to 0.75 seconds. During the next stage of training, self-performed trials, where both spouts moved within reach of the animal following stimulus presentation, were progressively introduced. An animal was considered trained when its detection performance across two or more sessions was above 80%.

### Behavioral monitoring

Data was collected from multiple sensors in the behavioral setup to measure different aspects of animal movement. Touch sensors using a grounding circuit on handles and lick spouts detected contact with the animal’s forepaws and tongue, respectively. A piezo sensor (1740, Adafruit LLC) below the animal’s trunk was used for monitoring body and hindlimb movements. Two webcams (C920 and B920, Logitech) were used for video recording of animal movements. Cameras were positioned to capture the animal’s face (side view) and the ventral surface of the body (ventral view).

### Widefield imaging

Widefield imaging was done as reported previously^23,32,81^ using an inverted tandem-lens macroscope in combination with an sCMOS camera (Edge 5.5, PCO) running at 30 fps. The top lens had a focal length of 105 mm (DC-Nikkor, Nikon) and the bottom lens 85 mm (85M-S, Rokinon). The field of view was 12.5 × 10.5 mm^2^ and the imaging resolution was 640 × 540 pixels after 4x spatial binning, resulting in a spatial resolution of ∼20 μm per pixel. To capture GCaMP fluorescence, a 525 nm band-pass filter (#86-963, Edmund optics) was placed in front of the camera. Using excitation light at two different wavelengths, we isolated Ca^2+^-dependent fluorescence and corrected for intrinsic signals (e.g., hemodynamic responses)^22,25^. Excitation light was projected on the cortical surface using a 495 nm long-pass dichroic mirror (T495lpxr, Chroma) placed between the two macro lenses. The excitation light was generated by a collimated blue LED (470 nm, M470L3, Thorlabs) and a collimated violet LED (405 nm, M405L3, Thorlabs) that were coupled into the same excitation path using a dichroic mirror (#87-063, Edmund optics). We alternated illumination between the two LEDs from frame to frame, resulting in one set of frames with blue and the other with violet excitation at 15 fps each. Excitation of GCaMP at 405 nm results in non-calcium dependent fluorescence^82^, allowing us to isolate the true calcium-dependent signal by rescaling and subtracting frames with violet illumination from the preceding frames with blue illumination. All subsequent analysis was based on this differential signal. The imaging data was then rigidly aligned to the Allen common coordinate framework (CCF), using four anatomical landmarks: the left, center and right points where anterior cortex meets the olfactory bulbs, and the medial point at the base of retrosplenial cortex. Retinotopic visual mapping experiments^31,83^ confirmed accurate CCF alignment and showed high correspondence between functionally identified visual areas and the CCF across PyN types (Fig. 1c).

### Two-photon imaging

We used a two-photon resonant scanning microscope (Moveable Objective Microscope, Sutter Instruments) for session-long, continuous image acquisition at 30.9 Hz. A 16X, 0.8 NA Nikon objective lens was used for single-plane imaging with a 512×512 pixels (575 µm x 575 µm) fields of view. Mode-locked illumination at 930 nm was delivered using a Ti:Sapphire laser (Ultra II, Coherent). Imaging frames were aligned with behavior control events synchronized acquisition of analog galvanometric and Bpod output signals. Depth of focal planes was 200-400 µm below the dura. Emission was collected using band-pass red (670/50 nm) and green (525/50 nm) filters (Chroma Technologies). MScan software (Sutter Instruments) was used for image acquisition. Recordings were performed in ALM (2.5 mm rostra and 1.5 mm lateral to bregma) or MM (1.5 mm anterior and 1 mm lateral to bregma) in randomized order across mice. Across imaging session, we selected planes that differed from those of prior sessions in order to maximize the number of unique neurons by.

Raw images were processed using the Suite2P package^84^ which performed motion correction, model-based ROI detection, correction for neuropil contamination, and spike deconvolution. Somatic and non-somatic (neuropil) ROI identification was performed through a combination of pre-trained classifier and manual curation. Somata with tdTomato expression were identified in a two-step process. First, potential green channel bleed-through was subtracted from the red channel using non-rigid regression with individual channels being divided into smaller blocks. Next, all sessions were then manually inspected to identify a conservative red fluorescence threshold, which was subsequently applied to all sessions. Analyses of neural activity were based on deconvolved values (“inferred spiking activity”). Since the deconvolved values do not represent absolute firing rates, we performed z-score normalization for each neuron before computing trial-averages across cells. The total number of recorded neurons per session was 396 ± 105 (mean ± standard deviation).

### Optogenetic inactivation

Photostimulation was performed using a 470 nm high-power LED (M470F3, Thorlabs) with a power density of 25 mW/mm^2^. Stimuli consisted of a square wave stimulus that ramped down in power for 200 milliseconds, to avoid an excitatory post-illumination rebound due to sudden release of inhibition^85^. To prevent animals’ visual detection of photostimulation, either through external leakage from light-insulated mating sleeves or transmission to the retina across the brain, an external LED with matching wavelength placed at the center of the animal’s visual field was flashed throughout the duration of every trial. Photoinhibition was performed in 20% of total trials and randomly interleaved between light-off trials. During each session, only bilateral frontal, parietal or visual cortex inhibition was performed. Once an animal was habituated and able to complete detection behavior trials with > 90% accuracy, optogenetic inactivation trials were introduced. During these initial sessions, optogenetic inhibition was performed from the beginning of the stimulus epoch until the end of the delay epoch. Additionally, we performed 0.5-second inhibition during four pre-defined epochs of the detection behavior trials: (1) first half of the stimulus, (2) second half of the stimulus, (3) delay, (4) response.

### Immunohistology, microscopy and image analysis

For a given animal, after all experiments were concluded, we performed transcardial perfusion with PBS followed by fixation with 4% PFA in 0.1 M PB. Brains were post-fixed in 4% PFA for an additional 12-18 hours at 4°. Prior to sectioning, brains were rinsed three times in PBS and embedded in 4% agarose-PBS. Slices 50 μm in thickness were made using a vibrating microtome (Leica, VT100S). Sections were then suspended in blocking solution (10% Normal Goat Serum and 0.1% Triton-X100 in 1X PBS) for 1 hour followed by overnight incubation at 4°C with the primary antibody. Next, sections were washed with PBS, incubated for 1 h at room temperature with the secondary antibody at 1:500 dilution. For visualization of GCaMP6s, we used primary goat polyclonal anti-GFP antibody (Abcam) and secondary donkey anti-goat Alexa Fluor 488 (Abcam). Sections were then dry-mounted on slides using Vectashield (Vector Labs, H1000) prior to imaging. No immunostaining was performed for the visualization of FusionRed or tdTomato. Imaging was performed using upright fluorescence macroscope and microscope (Olympus BX61). Images were acquired using Ocular Scientific Image Acquisition Software (Teledyne Imaging) and visualization and analysis were performed using ImageJ/FIJI software packages.

### Quantification cortex-wide gene expression

Quantification of cell counts across the dorsal cortex was performed using publicly available serial two-photon tomography (STPT) datasets. (http://www.brainimagelibrary.org/)^26^. Cre expression patterns for IT and PT neurons were characterized with data from eight mice, expressing either Cre-dependent GFP (PlexinD1-2A-CreER;Snap25-LSL-2A-EGFP) or tdTomato (Fezf2-2A-CreER;Ai14), respectively. Cell counting was then performed via automated soma detection, using a trained convolutional neural network^86^. STPT datasets were then registered to the Allen CCF v3 using the elastix toolbox^87^. To obtain the density of Cre-expressing neurons for individual cortical areas, we used the area outlines from the Allen CCF and computed the average sum of detected IT or PT neuron in each area, normalized by its surface area.

### Preprocessing of neural data

We first performed motion correction on each imaging frame, using a rigid-body image registration method implemented in the frequency domain^88^ that aligned each frame to the median over all frames in the first trial. To reduce the computational cost of subsequent analyses, we then computed the 200 highest-variance components using singular value decomposition (SVD). These components accounted for at least 95% of the total variance in each recording, whereas computing 500 components accounted for little additional variance (∼0.15%). SVD reduces the raw imaging data Y to a matrix of ‘spatial components’ U (of size pixels by components), ‘temporal components’ V^T^ (of size components by frames) and singular values S (of size components by components) to scale temporal components to the original data. The resulting decomposition has the form Y = USV^T^. All subsequent analysis in the time domain (such as the encoder and decoder models described below) were then performed on the product SV^T^ and the respective results were later multiplied with U, to recover results for the original pixel space. To avoid slow drift in the imaging data, SVT was high-pass filtered above 0.1 Hz using a zero-phase, second-order Butterworth filter.

To compute trial averages and perform choice decoder analysis (see below), imaging data in individual trials were aligned to the four trial periods, each marked by a specific event. This was required because the duration of different trial events was randomized to reduce temporal correlations, e.g. between trial initiation, the stimulus presentation and subsequent lick responses. The first period (Initiate) was aligned to the time when animal initiated a trial by touching the handles, the second (Stimulus) was aligned to the stimulus onset, the third (Delay) to the end of the stimulus sequence, and the fourth (Response) to the time when spouts were moved in to allow a lick response. After alignment, the total trial duration was 2 seconds and durations of respective trial episodes were 0.5 (Initiate), 1 (Stimulus), 0.2 (Delay), and 0.3 seconds (Response).

### Spatial clustering and classification

To obtain more interpretable spatial components and assess the dimensionality of cortical activity in different PyN types, we used semi-nonnegative matrix factorization (sNMF). As with SVD, sNMF also creates spatial and temporal components for each session and for each mouse but enforces spatial components to be strictly positive. The reason why temporal components were not also enforced to be non-negative is that our hemodynamic correction resulted in temporal dynamics that can be either positive or negative, relative to baseline. We used the LocaNMF toolbox by Saxena et al^33^ (https://github.com/ikinsella/locaNMF) to transform the spatial and temporal components U and SV^T^ into two corresponding matrices A and C, where A is a matrix of non-negative spatial components (of size pixels by components) and C the corresponding temporal components (of size components by frames). In addition to regular sNMF, the LocaNMF toolbox can be initialized with spatial constraints that are based on the Allen CCF. To obtain spatially restricted localized LocaNMF components, we constructed a map of larger seed regions by merging several smaller areas in the Allen CCF together (Fig. 2e). This region map is then used to enforce that each component in A is sparse outside the boundary of a given region. The amount of possible overlap between regions is specified by a localization threshold which specifies the percentage of a given component that is constrained to be inside a single region’s boundary. To obtain dense spatial components that were mostly driven by the local correlations between pixels and could also lie at the border between seed regions, we used a localization threshold of 50%. To obtain unconstrained sNMF spatial components, we also used the LocaNMF toolbox but only provided a single region that spanned the entire cortex. This resulted in cortex-wide components, similar to vanilla sNMF, while ensuring that all other analysis steps were done identically for sNMF and LocaNMF components. In both cases, we determined how many components in A and C were needed to explain 99% of the variance of Y (with Y=AC) after the initial SVD.

To compare spatial sNMF and LocaNMF components from different PyN types, we embedded them in a 2-dimensional space, using Uniform Manifold Approximation and Projection (UMAP) analysis (Fig. 2c,g). UMAP analysis was performed with the UMAP toolbox by McInnes et al.^35^ (https://github.com/lmcinnes/umap). For each recording, the first 20 spatial components in A (either from sNMF or region-constrained LocaNMF) were downsampled by a factor of 2, smoothed with a 2-D gaussian filter (5 × 5 pixels, 2 pixel standard deviation) and peak-normalized. Components from all recordings and animals were then combined into a larger matrix (of size pixels by components) and we used UMAP to project the first (pixel) dimension into two, maximally separating non-linear dimensions. Each point in the two dimensional space (Fig. 2c,g) therefore reflects a single component from one animal in a given imaging session.

The same UMAP approach was used for temporal sNMF and LocaNMF components. Before the UMAP projection, we first computed the trial-averaged and z-scored activity of each component to achieve temporal dynamics that are comparable across sessions and individual mice.

To identify PyN types based on individual spatial components (Fig. 2d, h), we performed a separate UMAP projection for each mouse. Each of these projections excluded all components from the test animal, ensuring that the UMAP projection was not shaped by potential noise patterns or other unknown features of the test components that could affected the classifier result. We then tested the first 20 components of each session of the test animal with 100 repetitions per component. In every repetition, 1000 components from each PyN type were randomly selected from the pre-computed UMAP space and we then assigned the PyN type of the test component based on the identity of its 10 nearest neighbors in UMAP space. For LocaNMF components, we performed the same procedure but additionally ensured to use an equal number of components from each seed region and PyN type to prevent PyN types with a larger number of components in a given region from biasing the classifier result. Classifier accuracy for each session (Fig. 2d,h) was then computed as the mean probability over all repetitions to accurately identify the PyN type.

To determine the size of PyN-predictive LocaNMF components, we selected all spatial components that achieved a classification accuracy of 99% percent or higher (all other components were assigned as non-specific) and thresholded each component above 0.2 to obtain a binary image. The size of each component was then computed as the square root of the sum of all pixels and converted to square millimeters.

### Linear encoding model

The linear encoding model was based on a combination of task- and movement-related variables, as described previously^32^. Each variable consisted of multiple regressors that were combined into a larger design matrix. Binary regressors contained a single pulse that signaled the occurrence of specific events, such as the stimulus onset, and additional regression copies that were shifted forward or backward in time to account for changes in cortical activity before or after the respective event. For auditory stimuli, the time-shifted copies spanned all frames from the onset of the auditory sequence until the end of the trial. Individual click sounds were also captured by an additional regressor set that spanned the 2 seconds after click onset. For motor events, like licking or whisking, the time-shifted copies spanned the frames from 1 s before until 2 s after each event. Lastly, for some variables, such as the previous choice, the time-shifted copies spanned the whole trial. Other variables were analog, such as measures from the piezo sensor or the pupil diameter, and also contained the 200 highest temporal components of video information from both cameras (using SVD as described above). This ensured that the model could account for animal movements and accurately isolate task-related activity. Movement and task variables were additionally decorrelated due to the variable durations of the initiation, stimulus and delay period. The model was fit using ridge regression to allow for similar contributions from different correlated variables. To determine the regularization penalty λ for each column of the widefield data, we used marginal maximum likelihood estimation (MLE)^89^. MLE expresses the encoding model as a Bayesian linear model and determines the ridge penalty λ by maximizing the marginal likelihood π(D | λ) of the model, given data D. This was done iteratively by testing different λ values to determine a global minimum for the negative log-likelihood −log π(D | λ). The main advantage of this approach is that λ can be determined without computationally expensive cross-validation procedures, resulting in a ∼50-fold decrease in required compute time on a regular work station. Moreover, the faster MLE approach allows adjusting λ values for individual widefield data components which results in higher cross-validated explained variance of the encoding model, compared to a regular cross-validation approach (Supp. Fig. 16).

### Variance analysis

Explained variance (cvR^2^) was obtained using 10-fold cross-validation. This was done by fitting the model weights to a continuous 90%-large section of the imaging data and then computing the explained variance in the remaining 10% of the data. The procedure was repeated for 10 times, while shifting the training and test data to ensure that each part of the recording was used in the test data in one of the folds. To assess unique explained variance by individual variables (ΔR^2^), we created reduced models in which all regressors of a specific variable were shuffled in time. Shuffling of each regressor was done within individual trials to account for a potential impact of very slow temporal correlations due the kinetics of the calcium indicator. The difference in explained variance between the full and the reduced model yielded the unique contribution ΔR^2^ of that model variable that could not be explained by other variables in the model. The same approach was used to compute unique contributions for groups of variables, i.e., ‘Movements’ and ‘Task’. Here, all variables that corresponded to a given group were shuffled at once.

### Decoding model

To predict animal’s left/right choices from widefield data, we trained logistic regression decoders with an L1 penalty on the temporal component matrix SV^T^ in each session. The L1 penalty was defined as the inverse of the number of observations in the test dataset during cross-validation, which yielded a good balance between the cross-validated prediction accuracy of the decoder and the number of non-zero model regressors. When decoding choice, we randomly removed trials until there was an equal amount of correct and incorrect trials where mice chose the left and the right side. By balancing left/right choice sides and correct/incorrect trials, we ensured that the decoder would not predict choices due to corresponding sensory information or would be influenced by potential side biases. The logistic regression model was implemented in Matlab using the ‘fitclinear’ function and run repeatedly for each time point in individual trials after re-aligning them to trial periods as described above. In each session, all decoder runs were performed with the same amount of trials (at least 250 trials) and we used 10-fold cross-validation to compute decoder accuracy at each time point in the trial. Beta weights were averaged from all models created during cross-validation and convolved with the spatial component matrix U to create cortical maps of the choice decoder weights.

### Receiver operating characteristic (ROC) analysis

We computed the area under the ROC curve (AUC) to quantify choice preference of single neurons obtained from two-photon imaging. AUCs were computed by comparing the mean neural activity during the stimulus and delay period in all trials with ipsilateral versus contralateral choices. AUC values denote the specificity of the neural activity to ipsi- or contralateral choices, with values below 0.5 signifying ipsilateral choice-selectivity and AUC values above 0.5 contralateral choice-selectivity. To identify statistically significant choice-selective neurons, AUC values were also computed for shuffled trial labels (randomly assigning ipsi- and contralateral choices across trials) for each neuron. This procedure was repeated 100 times to create a distribution of shuffled AUC values for each neuron. A neuron’s choice selectivity was then deemed significant if the probability of obtaining the actual AUC from the shuffled AUC distribution was less than 0.05.

## Code availability statement

Code for preprocessing (e.g., hemodynamic correction and dimensionality reduction) is available here: https://github.com/churchlandlab/WidefieldImager/tree/master/Analysis.

A cloud-based option for many of these steps is available at Neurocaas: http://neurocaas.org. To simplify the interaction with the online platform, a graphical user interface, dedicated to launching analysis and retrieving results from the NeuroCAAS platform is available here: (https://github.com/churchlandlab/wfield). Code for the ridge regression used in the encoding model is available on the lab’s GitHub page: https://github.com/churchlandlab/ridgeModel.

## Data availability statement

We will make preprocessed datasets from all animals available at time of publication. As in previous publications, we will also include a “readme” statement that explains how the data are stored. An example dataset has already been shared (DOI: 10.5281/zenodo.5834513) on Zenodo to demonstrate the feasibility of this pipeline: https://zenodo.org/record/5834513#.Yd3jHViZPX8.

**Supplementary Table 1:** Overview of different model variables.

| Variable name | Description | Variable type | Category |
| --- | --- | --- | --- |
| Hindlimb | Piezo sensor below the animal | Analog + Event kernel | Movement |
| Handles (Left / Right) | Touch events from handle sensors | Event kernel | Movement |
| Licks (Left / Right) | Lick events from spout sensors | Event kernel | Movement |
| Pupil | Pupil diameter, extracted from face camera | Analog + Event kernel | Movement |
| Nose | Nose movements, extracted from face camera | Analog + Event kernel | Movement |
| Whisking | Whisker movements, extracted from face camera | Analog + Event kernel | Movement |
| Body | Average motion energy across all body camera pixels | Analog + Event kernel | Movement |
| Video | Video dimensions from both cameras (SVD) | Analog | Movement |
| Video ME | Video dimensions from motion energy in both cameras | Analog | Movement |
| Choice (Left / Right) | All frames in either a left- or a rightward choice trial | Event kernel | Task |
| Previous choice | Every trial after a leftward choice trial | Event kernel | Task |
| Previous modality | Every trial after a visual trial | Event kernel | Task |
| Previous success | Every trial after a successful trial | Event kernel | Task |
| Success | All successful trials | Event kernel | Task |
| Water given | All frames after a water reward was given | Event kernel | Task |
| Auditory stimulus (Left / Right) | All frames after a left- or rightward auditory stimulus | Event kernel | Task |
| Visual stimulus (Left / Right) | All frames after a left- or rightward visual stimulus | Event kernel | Task |

**Supplementary Table 2:**
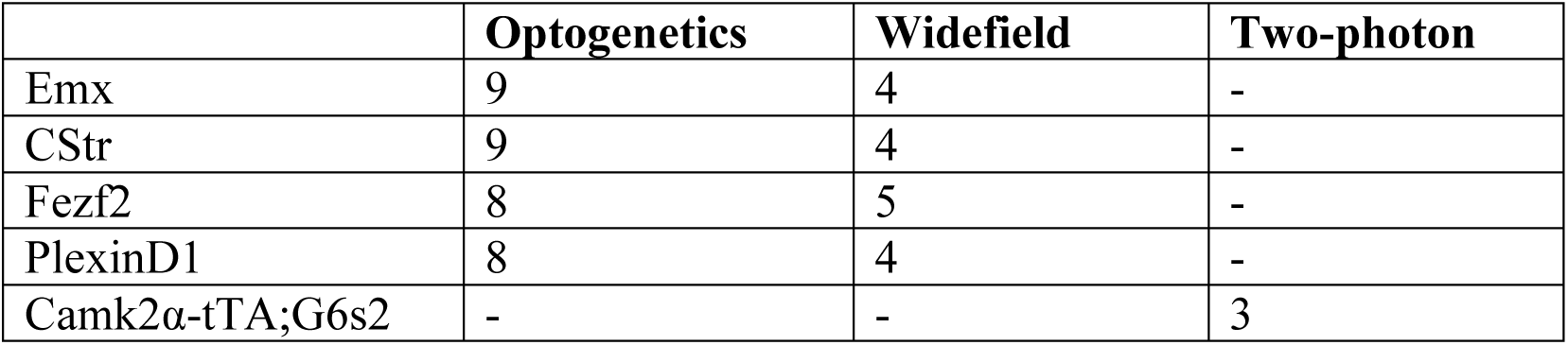
Number of mice included for each experiment.

|  | Optogenetics | Widefield | Two-photon |
| --- | --- | --- | --- |
| Emx | 9 | 4 | - |
| CStr | 9 | 4 | - |
| Fezf2 | 8 | 5 | - |
| PlexinD1 | 8 | 4 | - |
| Camk2α-tTA;G6s2 | - | - | 3 |

**Supplementary Table 3:**
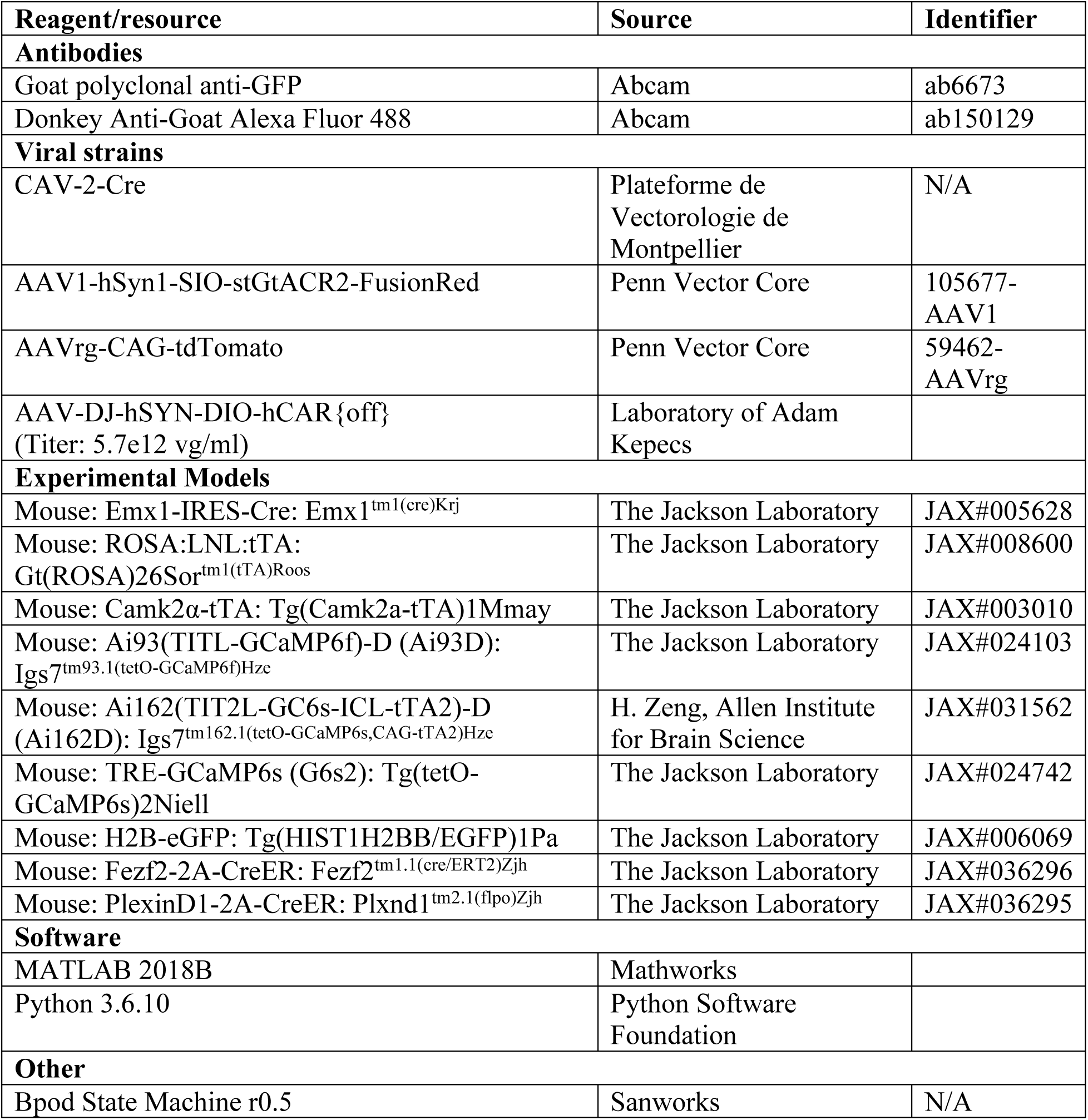
Resources.

## References

1. Harris, K. D. & Mrsic-Flogel, T. D. Cortical connectivity and sensory coding. Nature 503, 51–58 (2013).

2. Pfeffer, C. K., Xue, M., He, M., Huang, Z. J. & Scanziani, M. Inhibition of Inhibition in Visual Cortex: The Logic of Connections Between Molecularly Distinct Interneurons. Nat Neurosci 16, 1068–1076 (2013).

3. Keller, A. J. et al. A Disinhibitory Circuit for Contextual Modulation in Primary Visual Cortex. Neuron 108, 1181-1193.e8 (2020).

4. Tremblay, R., Lee, S. & Rudy, B. GABAergic Interneurons in the Neocortex: From Cellular Properties to Circuits. Neuron 91, 260–292 (2016).

5. Cardin, J. A. et al. Driving fast-spiking cells induces gamma rhythm and controls sensory responses. Nature 459, 663–667 (2009).

6. Veit, J., Hakim, R., Jadi, M. P., Sejnowski, T. J. & Adesnik, H. Cortical gamma band synchronization through somatostatin interneurons. Nature Neuroscience 20, 951–959 (2017).

7. Sohal, V. S., Zhang, F., Yizhar, O. & Deisseroth, K. Parvalbumin neurons and gamma rhythms enhance cortical circuit performance. Nature 459, 698–702 (2009).

8. Reimer, J. et al. Pupil fluctuations track fast switching of cortical states during quiet wakefulness. Neuron 84, 355–362 (2014).

9. Fu, Y. et al. A Cortical Circuit for Gain Control by Behavioral State. Cell 156, 1139–1152 (2014).

10. Zhou, M. et al. Scaling down of balanced excitation and inhibition by active behavioral states in auditory cortex. Nature Neuroscience 17, 841–850 (2014).

11. Polack, P.-O., Friedman, J. & Golshani, P. Cellular mechanisms of brain state–dependent gain modulation in visual cortex. Nature Neuroscience 16, 1331–1339 (2013).

12. Tasic, B. et al. Shared and distinct transcriptomic cell types across neocortical areas. Nature 563, 72–78 (2018).

13. Economo, M. N. et al. Distinct descending motor cortex pathways and their roles in movement. Nature 563, 79–84 (2018).

14. Chen, J. L., Carta, S., Soldado-Magraner, J., Schneider, B. L. & Helmchen, F. Behaviour-dependent recruitment of long-range projection neurons in somatosensory cortex. Nature 499, 336–340 (2013).

15. Glickfeld, L. L., Andermann, M. L., Bonin, V. & Reid, R. C. Cortico-cortical projections in mouse visual cortex are functionally target specific. Nat Neurosci 16, 219–226 (2013).

16. Kim, T. et al. Cortically projecting basal forebrain parvalbumin neurons regulate cortical gamma band oscillations. PNAS 112, 3535–3540 (2015).

17. Harris, K. D. & Shepherd, G. M. G. The neocortical circuit: themes and variations. Nature Neuroscience 18, 170–181 (2015).

18. Takahashi, N. et al. Active dendritic currents gate descending cortical outputs in perception. Nature Neuroscience 1–9 (2020) doi:10.1038/s41593-020-0677-8.

19. Tang, L. & Higley, M. J. Layer 5 Circuits in V1 Differentially Control Visuomotor Behavior. Neuron 105, 346-354.e5 (2020).

20. Li, N., Chen, T.-W., Guo, Z. V., Gerfen, C. R. & Svoboda, K. A motor cortex circuit for motor planning and movement. Nature 519, 51–56 (2015).

21. Chen, T.-W., Li, N., Daie, K. & Svoboda, K. A Map of Anticipatory Activity in Mouse Motor Cortex. Neuron 94, 866-879.e4 (2017).

22. Wekselblatt, J. B., Flister, E. D., Piscopo, D. M. & Niell, C. M. Large-scale imaging of cortical dynamics during sensory perception and behavior. Journal of Neurophysiology 115, 2852–2866 (2016).

23. Couto, J. et al. Chronic, cortex-wide imaging of specific cell populations during behavior. Nat Protoc 1–25 (2021) doi:10.1038/s41596-021-00527-z.

24. Cardin, J. A., Crair, M. C. & Higley, M. J. Mesoscopic Imaging: Shining a Wide Light on Large-Scale Neural Dynamics. Neuron 108, 33–43 (2020).

25. Allen, W. E. et al. Global Representations of Goal-Directed Behavior in Distinct Cell Types of Mouse Neocortex. Neuron 94, 891-907.e6 (2017).

26. Matho, K. S. et al. Genetic dissection of the glutamatergic neuron system in cerebral cortex. Nature 598, 182–187 (2021).

27. Gerfen, C. R., Paletzki, R. & Heintz, N. GENSAT BAC cre-recombinase driver lines to study the functional organization of cerebral cortical and basal ganglia circuits. Neuron 80, 1368–1383 (2013).

28. Harris, J. A. et al. Anatomical characterization of Cre driver mice for neural circuit mapping and manipulation. Front Neural Circuits 8, 76 (2014).

29. Daigle, T. L. et al. A Suite of Transgenic Driver and Reporter Mouse Lines with Enhanced Brain-Cell-Type Targeting and Functionality. Cell 174, 465-480.e22 (2018).

30. Wang, Q. et al. The Allen Mouse Brain Common Coordinate Framework: A 3D Reference Atlas. Cell 181, 936-953.e20 (2020).

31. Garrett, M. E., Nauhaus, I., Marshel, J. H. & Callaway, E. M. Topography and Areal Organization of Mouse Visual Cortex. J. Neurosci. 34, 12587–12600 (2014).

32. Musall, S., Kaufman, M. T., Juavinett, A. L., Gluf, S. & Churchland, A. K. Single-trial neural dynamics are dominated by richly varied movements. Nat Neurosci 22, 1677–1686 (2019).

33. Saxena, S. et al. Localized semi-nonnegative matrix factorization (LocaNMF) of widefield calcium imaging data. PLOS Computational Biology 16, e1007791 (2020).

34. Condylis, C. et al. Dense functional and molecular readout of a circuit hub in sensory cortex. Science 375, eabl5981 (2022).

35. McInnes, L., Healy, J. & Melville, J. UMAP: Uniform Manifold Approximation and Projection for Dimension Reduction. arXiv:1802.03426 [cs, stat] (2018).

36. Brunton, B. W., Botvinick, M. M. & Brody, C. D. Rats and Humans Can Optimally Accumulate Evidence for Decision-Making. Science 340, 95–98 (2013).

37. Salkoff, D. B., Zagha, E., McCarthy, E. & McCormick, D. A. Movement and Performance Explain Widespread Cortical Activity in a Visual Detection Task. Cereb Cortex 30, 421–437 (2020).

38. Orsolic, I., Rio, M., Mrsic-Flogel, T. D. & Znamenskiy, P. Mesoscale cortical dynamics reflect the interaction of sensory evidence and temporal expectation during perceptual decision-making. Neuron 109, 1861-1875.e10 (2021).

39. Steinmetz, N. A., Zatka-Haas, P., Carandini, M. & Harris, K. D. Distributed coding of choice, action and engagement across the mouse brain. Nature 576, 266–273 (2019).

40. Bergmann, R. et al. Predicting behavior from eye movement and whisking asymmetry. 2021.02.11.430785 Preprint at https://doi.org/10.1101/2021.02.11.430785 (2021).

41. Esmaeili, V. et al. Rapid suppression and sustained activation of distinct cortical regions for a delayed sensory-triggered motor response. Neuron 109, 2183-2201.e9 (2021).

42. Hanks, T. D. et al. Distinct relationships of parietal and prefrontal cortices to evidence accumulation. Nature 520, 220–223 (2015).

43. Goard, M. J., Pho, G. N., Woodson, J. & Sur, M. Distinct roles of visual, parietal, and frontal motor cortices in memory-guided sensorimotor decisions. eLife 5, e13764 (2016).

44. Peters, A. J., Fabre, J. M. J., Steinmetz, N. A., Harris, K. D. & Carandini, M. Striatal activity topographically reflects cortical activity. Nature 1–6 (2021) doi:10.1038/s41586-020-03166-8.

45. Oh, S. W. et al. A mesoscale connectome of the mouse brain. Nature 508, 207–214 (2014).

46. Mahn, M. et al. High-efficiency optogenetic silencing with soma-targeted anion-conducting channelrhodopsins. Nature Communications 9, 4125 (2018).

47. Li, S.-J., Vaughan, A., Sturgill, J. F. & Kepecs, A. A Viral Receptor Complementation Strategy to Overcome CAV-2 Tropism for Efficient Retrograde Targeting of Neurons. Neuron 98, 905-917.e5 (2018).

48. Znamenskiy, P. & Zador, A. M. Corticostriatal neurones in auditory cortex drive decisions during auditory discrimination. Nature 497, 482–485 (2013).

49. Guo, Z. V. et al. Flow of Cortical Activity Underlying a Tactile Decision in Mice. Neuron 81, 179–194 (2014).

50. MacDowell, C. J. & Buschman, T. J. Low-Dimensional Spatiotemporal Dynamics Underlie Cortex-wide Neural Activity. Current Biology 30, 2665-2680.e8 (2020).

51. Stringer, C. et al. Spontaneous behaviors drive multidimensional, brainwide activity. Science 364, eaav7893 (2019).

52. Logothetis, N. K. et al. Hippocampal-cortical interaction during periods of subcortical silence. Nature 491, 547–553 (2012).

53. Rogers, B. P., Morgan, V. L., Newton, A. T. & Gore, J. C. Assessing Functional Connectivity in the Human Brain by FMRI. Magn Reson Imaging 25, 1347–1357 (2007).

54. Vanni, M. P. & Murphy, T. H. Mesoscale Transcranial Spontaneous Activity Mapping in GCaMP3 Transgenic Mice Reveals Extensive Reciprocal Connections between Areas of Somatomotor Cortex. J. Neurosci. 34, 15931–15946 (2014).

55. Vanni, M. P., Chan, A. W., Balbi, M., Silasi, G. & Murphy, T. H. Mesoscale Mapping of Mouse Cortex Reveals Frequency-Dependent Cycling between Distinct Macroscale Functional Modules. J. Neurosci. 37, 7513–7533 (2017).

56. Macé, É. et al. Whole-Brain Functional Ultrasound Imaging Reveals Brain Modules for Visuomotor Integration. Neuron 100, 1241-1251.e7 (2018).

57. Matsui, T., Murakami, T. & Ohki, K. Transient neuronal coactivations embedded in globally propagating waves underlie resting-state functional connectivity. PNAS 113, 6556–6561 (2016).

58. Huang, L. et al. BRICseq Bridges Brain-wide Interregional Connectivity to Neural Activity and Gene Expression in Single Animals. Cell 182, 177-188.e27 (2020).

59. Mohajerani, M. H. et al. Spontaneous cortical activity alternates between motifs defined by regional axonal projections. Nat Neurosci 16, 1426–1435 (2013).

60. Lohani, S. et al. Dual color mesoscopic imaging reveals spatiotemporally heterogeneous coordination of cholinergic and neocortical activity. 2020.12.09.418632 https://www.biorxiv.org/content/10.1101/2020.12.09.418632v1 (2020) doi:10.1101/2020.12.09.418632.

61. Gilad, A. & Helmchen, F. Spatiotemporal refinement of signal flow through association cortex during learning. Nature Communications 11, 1–14 (2020).

62. Gallero-Salas, Y. et al. Sensory and Behavioral Components of Neocortical Signal Flow in Discrimination Tasks with Short-Term Memory. Neuron 109, 135-148.e6 (2021).

63. Park, J. M. et al. Deep and superficial layers of the primary somatosensory cortex are critical for whisker-based texture discrimination in mice. bioRxiv 2020.08.12.245381 (2020) doi:10.1101/2020.08.12.245381.

64. Gilad, A., Gallero-Salas, Y., Groos, D. & Helmchen, F. Behavioral Strategy Determines Frontal or Posterior Location of Short-Term Memory in Neocortex. Neuron 99, 814-828.e7 (2018).

65. Jun, E. J. et al. Causal role for the primate superior colliculus in the computation of evidence for perceptual decisions. Nat Neurosci (2021) doi:10.1038/s41593-021-00878-6.

66. Wang, L., McAlonan, K., Goldstein, S., Gerfen, C. R. & Krauzlis, R. J. A Causal Role for Mouse Superior Colliculus in Visual Perceptual Decision-Making. J. Neurosci. 40, 3768– 3782 (2020).

67. Felsen, G. & Mainen, Z. F. Neural substrates of sensory-guided locomotor decisions in the rat superior colliculus. Neuron 60, 137–148 (2008).

68. Duan, C. A. et al. A cortico-collicular pathway for motor planning in a memory-dependent perceptual decision task. Nature Communications 12, 2727 (2021).

69. Erlich, J. C., Brunton, B. W., Duan, C. A., Hanks, T. D. & Brody, C. D. Distinct effects of prefrontal and parietal cortex inactivations on an accumulation of evidence task in the rat. eLife Sciences 4, e05457 (2015).

70. Raposo, D., Kaufman, M. T. & Churchland, A. K. A category-free neural population supports evolving demands during decision-making. Nat Neurosci 17, 1784–1792 (2014).

71. Harvey, C. D., Coen, P. & Tank, D. W. Choice-specific sequences in parietal cortex during a virtual-navigation decision task. Nature 484, 62–68 (2012).

72. Pinto, L. et al. Task-Dependent Changes in the Large-Scale Dynamics and Necessity of Cortical Regions. Neuron 104, 810-824.e9 (2019).

73. Licata, A. M. et al. Posterior Parietal Cortex Guides Visual Decisions in Rats. J. Neurosci. 37, 4954–4966 (2017).

74. Zatka-Haas, P., Steinmetz, N. A., Carandini, M. & Harris, K. D. Sensory coding and the causal impact of mouse cortex in a visual decision. eLife 10, e63163 (2021).

75. Kim, J.-H., Ma, D.-H., Jung, E., Choi, I. & Lee, S.-H. Gated feedforward inhibition in the frontal cortex releases goal-directed action. Nat Neurosci 24, 1452–1464 (2021).

76. Wu, Z. et al. Context-Dependent Decision Making in a Premotor Circuit. Neuron 106, 316-328.e6 (2020).

77. Coen, P., Sit, T. P. H., Wells, M. J., Carandini, M. & Harris, K. D. Mouse frontal cortex mediates additive multisensory decisions. 2021.04.26.441250 https://www.biorxiv.org/content/10.1101/2021.04.26.441250v2 (2021) doi:10.1101/2021.04.26.441250.

78. Park, J., Phillips, J. W., Martin, K. A., Hantman, A. W. & Dudman, J. T. Descending neocortical output critical for skilled forelimb movements is distributed across projection cell classes. 772517 https://www.biorxiv.org/content/10.1101/772517v3 (2021) doi:10.1101/772517.

79. Callaway, E. M. et al. A multimodal cell census and atlas of the mammalian primary motor cortex. Nature 598, 86–102 (2021).

80. Laboulaye, M. A., Duan, X., Qiao, M., Whitney, I. E. & Sanes, J. R. Mapping Transgene Insertion Sites Reveals Complex Interactions Between Mouse Transgenes and Neighboring Endogenous Genes. Frontiers in Molecular Neuroscience 11, 385 (2018).

81. Ratzlaff, E. H. & Grinvald, A. A tandem-lens epifluorescence macroscope: hundred-fold brightness advantage for wide-field imaging. J. Neurosci. Methods 36, 127–137 (1991).

82. Lerner, T. N. et al. Intact-Brain Analyses Reveal Distinct Information Carried by SNc Dopamine Subcircuits. Cell 162, 635–647 (2015).

83. Marshel, J. H., Garrett, M. E., Nauhaus, I. & Callaway, E. M. Functional Specialization of Seven Mouse Visual Cortical Areas. Neuron 72, 1040–1054 (2011).

84. Pachitariu, M. et al. Suite2p: beyond 10,000 neurons with standard two-photon microscopy. bioRxiv 061507 (2016) doi:10.1101/061507.

85. Chuong, A. S. et al. Noninvasive optical inhibition with a red-shifted microbial rhodopsin. Nat Neurosci 17, 1123–1129 (2014).

86. Kim, Y. et al. Brain-wide Maps Reveal Stereotyped Cell-Type-Based Cortical Architecture and Subcortical Sexual Dimorphism. Cell 171, 456-469.e22 (2017).

87. Klein, S., Staring, M., Murphy, K., Viergever, M. A. & Pluim, J. P. W. elastix: A Toolbox for Intensity-Based Medical Image Registration. IEEE Transactions on Medical Imaging 29, 196–205 (2010).

88. Reddy, B. S. & Chatterji, B. N. An FFT-based technique for translation, rotation, and scale-invariant image registration. IEEE Trans Image Process 5, 1266–1271 (1996).

89. Karabatsos, G. Marginal maximum likelihood estimation methods for the tuning parameters of ridge, power ridge, and generalized ridge regression. Communications in Statistics - Simulation and Computation (2017).

