## Supplementary figures for "Pyramidal cell types drive functionally distinct cortical activity patterns during decision-making"

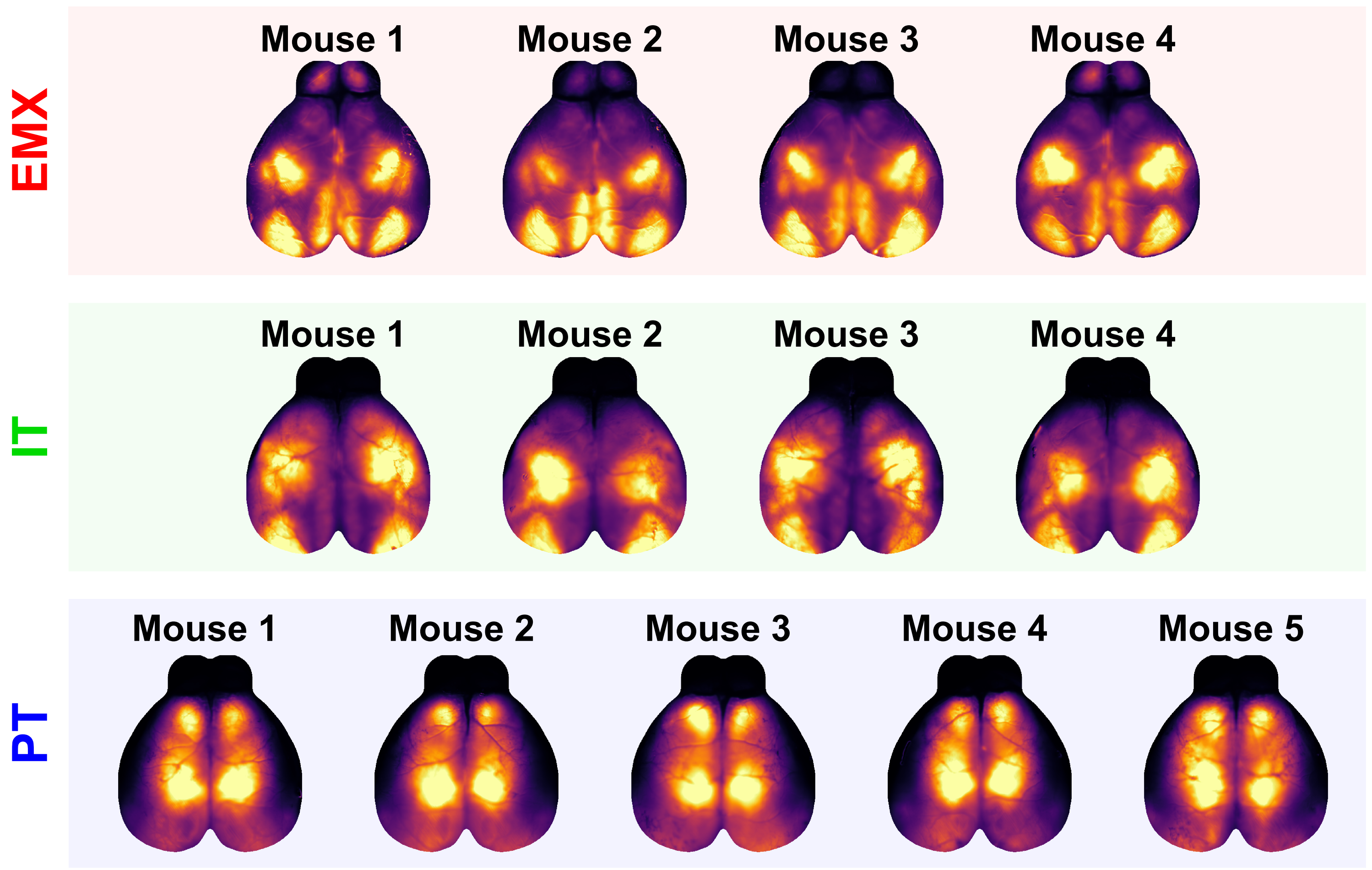


**Supplementary Fig. 1. Cortical maps of total variance for individual mice**
Maps of variance over all frames for individual mice in each PyN type group. Colors are normalized between zero and the 95th percentile for each animal. Distinct variance patterns for each PyN type were largely conserved across individual mice.

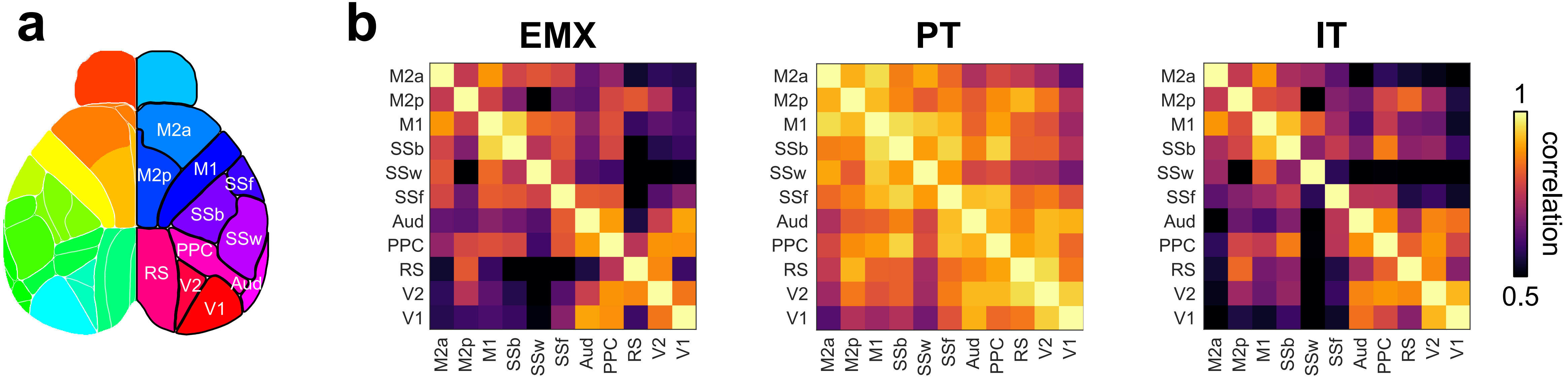


**Supplementary Fig. 2. PyN-specific correlation patterns between cortical regions**
**a)** Map of cortical regions, used for correlation analysis. V1 = primary visual cortex, V2 = secondary visual cortex, RS = retrosplenial cortex, Aud = auditory cortex, PPC = posterior parietal cortex, SSw = somatosensory whisker area, SSb = somatosensory body area, SSf = somatosensory face area, M1 = primary motor cortex, M2p = posterior secondary motor cortex, M2a = anterior secondary motor cortex. **b)** Correlations between cortical regions in EMX, PT and IT neurons averaged over all sessions and mice. Inter-region correlations were comparable between EMX and IT neurons but overall increased for PT neurons.

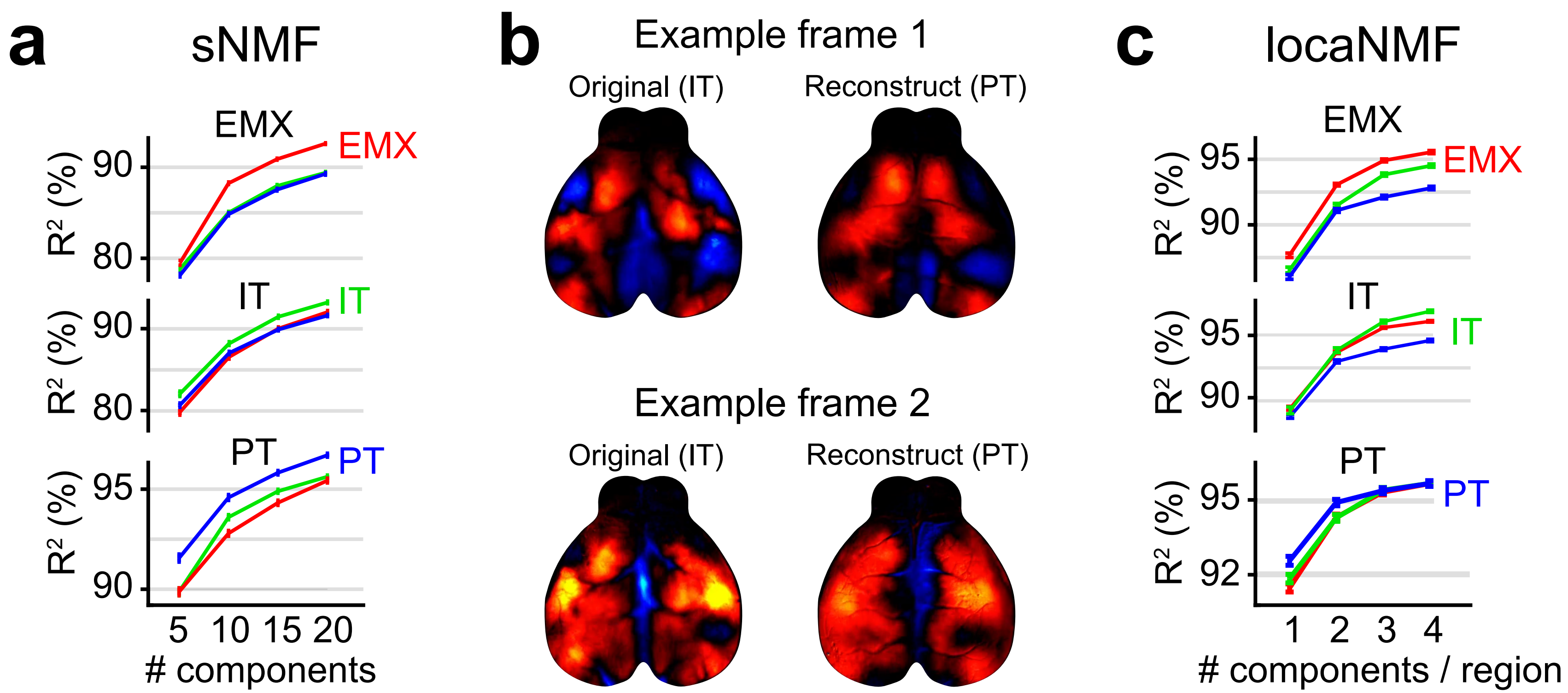


**Supplementary Fig. 3. Overlap of low-dimensional space and examples of single-frame reconstructions**

**a)** $R^2$ of EMX, IT and PT reconstructions (top to bottom panels), using components from different PyN types (red, green, and blue traces). For within-group reconstructions, only components from other mice were used. **b)** Single-frame reconstructions of IT data, using PT components. IT imaging data (original) was projected onto PT components to assess if they would be applicable to capture IT variance. While the reconstruction captured a large fraction of variance (~93%, Supplementary Video 4), comparing individual frames showed that PT components did not recreate more fine-grained spatial features of the IT data. **c)** $R^2$ of EMX, IT and PT reconstructions, using locaNMF components from different PyN types (formatting as in panel a)). Shown are results for different number of components per region, using 24 regions in total. The minum number of components was therefore 24 (1 component per region) and the maximum 96 (4 components per region).

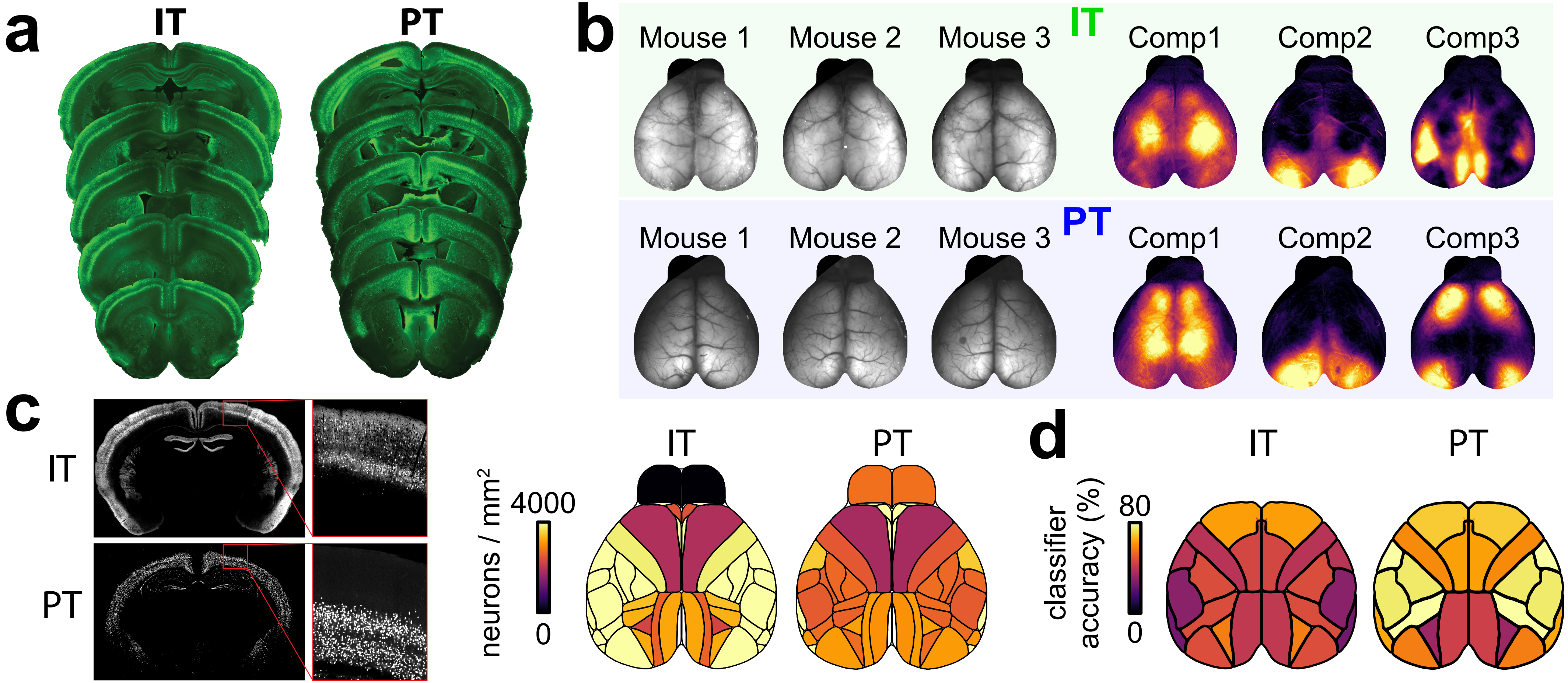


**Supplementary Fig. 4. Cortex-wide expression patterns of PT and IT neurons**

**a)** Brain slices from IT and PT neurons show robust cortex-wide expression of GCaMP6. **b)** Left: Raw fluorescence from widefield imaging of 3 different PT and IT mice. In both lines, we obtained strong fluorescence throughout the cortex, although minor fluctuations in brightness were visible across regions. Right: Example sNMF components from an invidiual PT/IT mouse (mouse #3 for both lines). sNMF components did not strongly reflect differences in raw fluoresence across cortex. We also observed no clear relationship between fluoresence patterns to total variance (Fig. 1D) or ongoing activity patterns (Supp. Movies 2, 3). **c)** Left: Example brain slices from IT- and PT-Cre mice to quantify the density of Cre-expressing neurons in each line. Blow-up shows a magnified region in cortex with individual somata. Right: Expression density was largely even across dorsal cortex with higher density of IT neurons in lateral regions and no expression in the olfactory bulb. Density was slightly reduced in M2 for both lines. **d)** Map of PyN type decoding accuracy with locaNMF components for different cortical regions. Decoding accuracy was high across cortical regions and we found no clear relation betwen expression patterns and regions with particularly high locaNMF decoding accuracy. Olfacory bulb was omitted from the analysis, due to the lack of strong fluorescence signals.

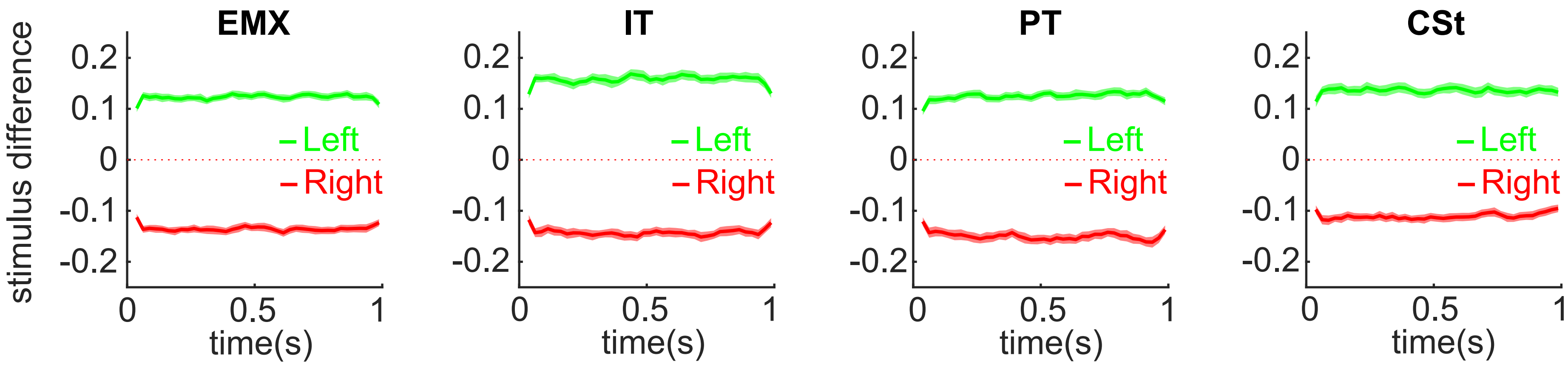


**Supplementary Fig. 5. Mice in all PyN groups integrate sensory information throughout the stimulus period**
Shown is the normalized difference between auditory clicks on the left or right side, when animals succesfully responded to the left (green) or the right (red). Binsize is 50 ms. Positive numbers indicate a higher probability of observing a leftward click sound, negative numbers indicate more clicks on the right. In all mice, the probability of observing more stimuli on the correct side is consistently higher throughout the stimulus period. This shows that mice integrate sensory evidence from the entire stimulus period and auditory clicks equally influence animal decisions, regardless of whether they occur early or late in the stimulus sequence.

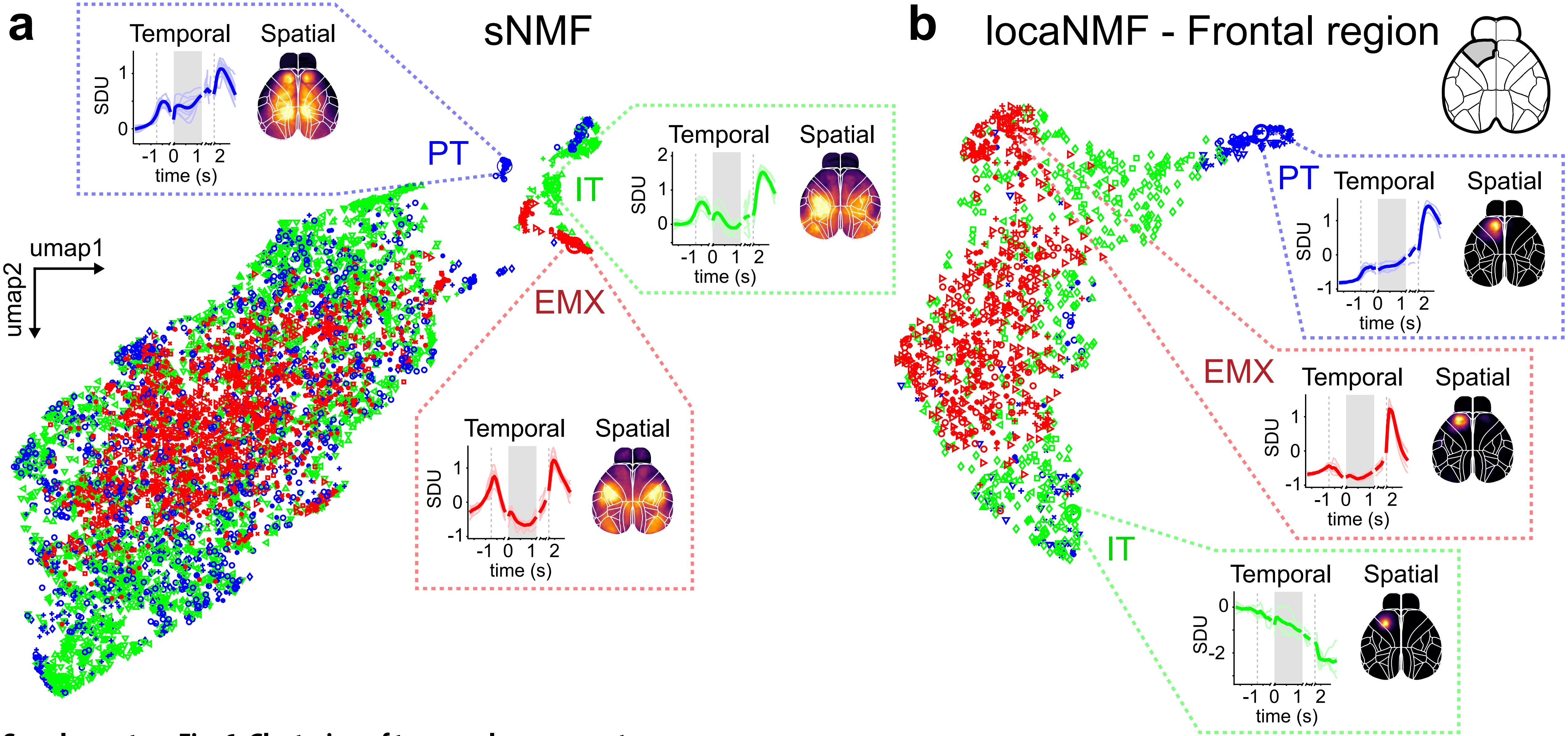


**Supplementary Fig. 6. Clustering of temporal components**

**a)** UMAP embedding of temporal sNMF components for EMX (red), IT (green) and PT (blue) mice. Clustering for cell types is weaker as with spatial components (Fig. 2c) but clearly visible, suggesting that sNMF components are both spatially and functionally distinct. Insets show 10 example traces of trial-averaged activity from cell-type specific clusters (left, bold line shows the mean) and an example of a corresponding spatial component (right). **b)** UMAP embedding of temporal locaNMF components from left frontal cortex. Conventions as in a). Temporal locaNMF components also show cell- type-specific clustering, revealing task-specific dynamics (inset, left). Spatial locaNMF components also show separate shapes for each cell type (inset, right).

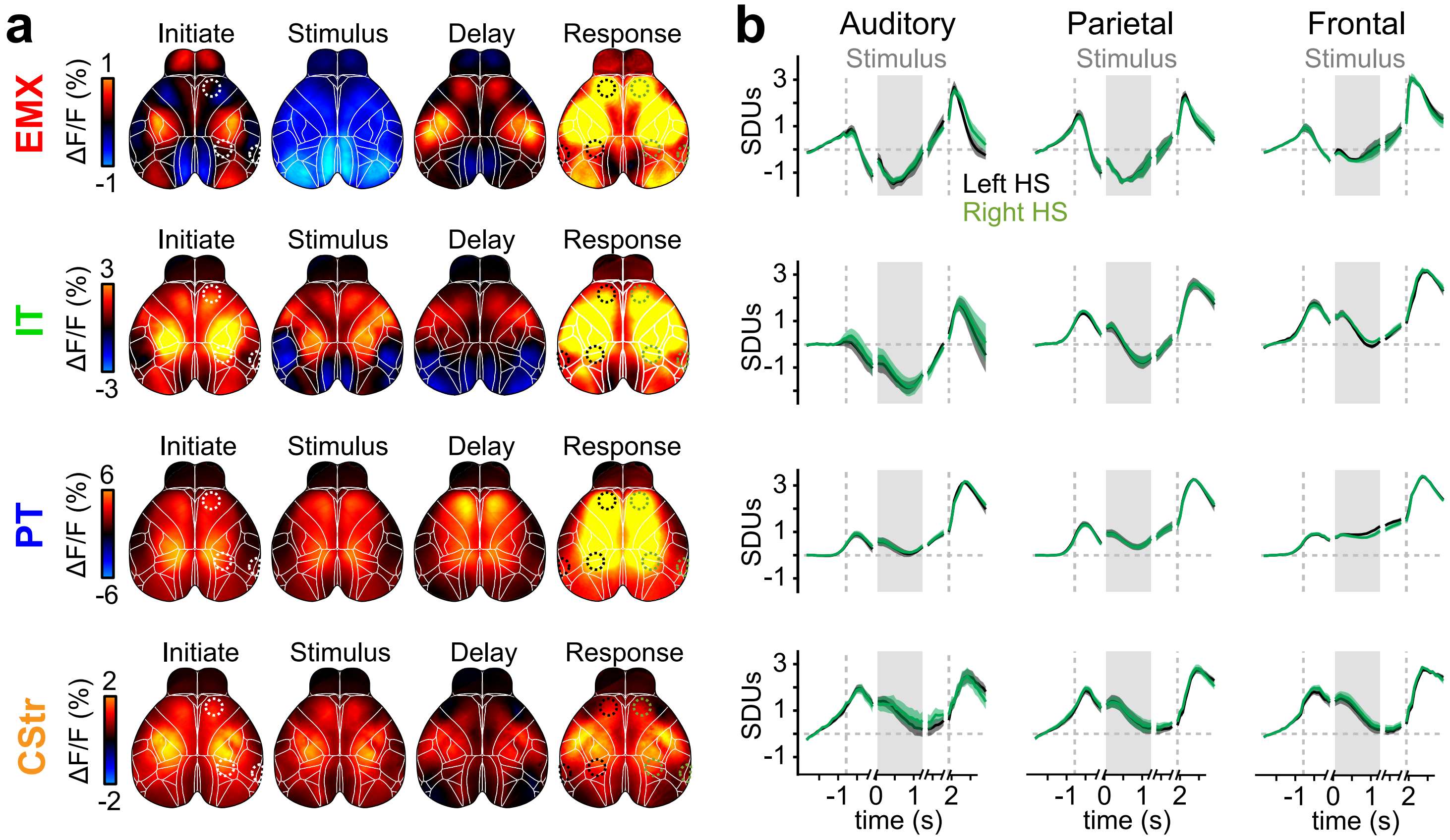


**Supplementary Fig. 7. Symmetric bilateral activation during decision-making**

**a)** Trial-averaged response maps for all correct, leftward trials across different PyN types. Cortical maps are as shown in Fig. 3a and Fig. 7c. **b)** Average activity in auditory, parietal, and frontal cortex on the left (black) and right hemisphere (green), which are contra- and ipsilateral to the chosen side, respectively. In all PyN types, different trial events, such as initiation, sensory stimulation and animal responses increased neural activity. However, surprisingly few differences were seen between cortical hemisspheres. To resolve differences in inter-hemispheric activation for left- versus rightward choices we therefore employed the choice decoder analysis (Fig. 6 and 7). Note that low or negative weights from the choice decoder (as seen for IT and CStr neurons in frontal cortex) do not reflect a lack of choice-related activity activity but are rather based on small differences in the activation of hemispheres that are either ipsi- or contralateral to the chosen side.

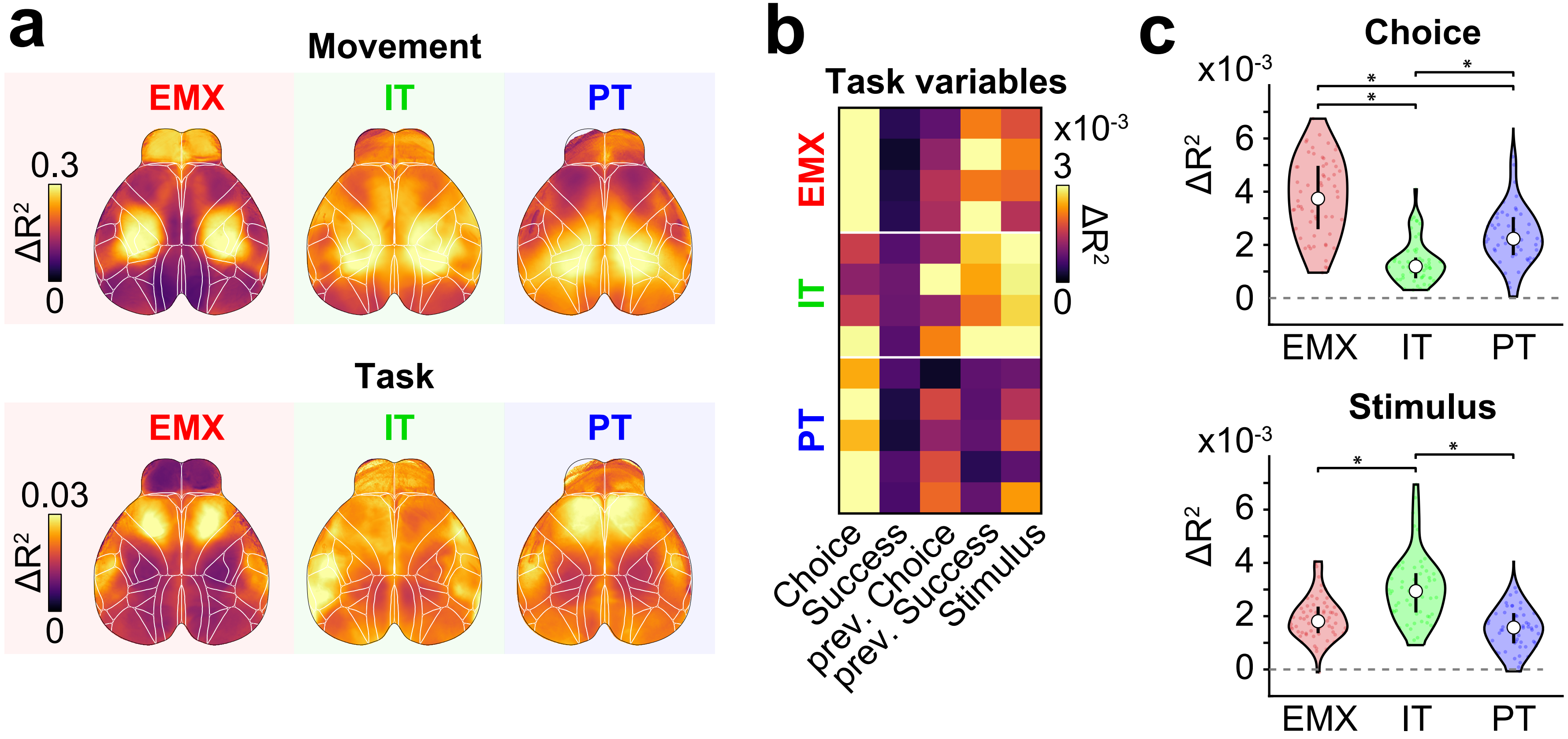


**Supplementary Fig. 8. Unique explained variance of model variables**

**a)** To isolate unique contributions from movement or task variables, we computed averaged maps of the loss in predicted variance ($\Delta R^2$) by removing either group of variables from the full model. This allowed us to separately examine their respective impact on cortex-wide activity by determining, for each PyN type, where in the cortex predictive power was lost. While movement $\Delta R^2$ patterns were comparable across PyN types (top row), PyN-type-specific differences were uncovered when removing task variables: $\Delta R^2$ was highest in frontal cortex of EMX and PT mice, but more diffuse in IT mice with the highest $\Delta R^2$ in auditory cortex (bottom row). Note differences in scale between two rows. **b)** Examination of $\Delta R^2$ for individual task variables further suggest distinct roles for each PyN type. Here, the 'choice' variable had the highest contributions in PT neurons but was overall weaker in IT neurons. Conversely, contributions from other task variables were higher in IT neurons. This dichotomy was not observed in EMX neurons, indicating that IT and PT neurons might have different functional roles that cannot be resolved without PyN-type specific measurements. Each row represent a mouse. **c)** Comparison of $\Delta R^2$ for choice (top) and simulus variables (bottom) between PyN types. IT mice had significantly lower $\Delta R^2$ for choice but higher $\Delta R^2$ for the stimulus as EMX or PT mice. Note that lower $\Delta R^2$ for choice in IT mice does not imply a lack of involvement in decision formation but rather that their population activity does not clearly differ for left versus right choices. Dots indicate individual sessions. Stars indicate significant differences across sessions (*t*-test, $p<0.05$, bonferroni-corrected).

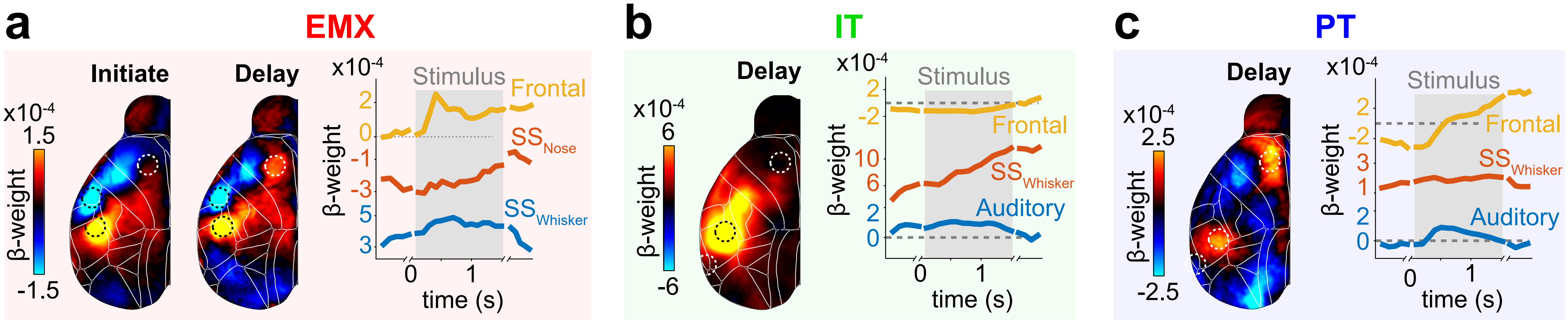


**Supplementary Fig. 9. Choice-related activity in somatosensory cortex**

**a)** Averaged choice kernel maps for EMX mice during the initiation and delay period. Dashed circles show location of somatosensory whisker ($SS_{Whisker}$), somatosensory nose ($SS_{Nose}$), and frontal cortex. $SS_{Whisker}$ (blue trace) and $SS_{Nose}$ (red trace) were constantly positive or negative, respectively, even during the initiation period. In contrast to frontal cortex (yellow trace), both areas were only weakly modulation by the stimulus onset (gray box). **b)** Choice kernel maps for IT mice during the delay period. Dashed circles show location of auditory, $SS_{Whisker}$, and frontal cortex. Choice-related activity in $SS_{Whisker}$ (red trace) increased over the course of the trial. No choice-related modulation was apparent in frontal cortex. **c)** Choice kernel maps for PT mice during the delay period. Conventions as in b). Choice-related activity strongly increased in frontal crontal cortex after stimulus onset and was weaker in other cortical areas.

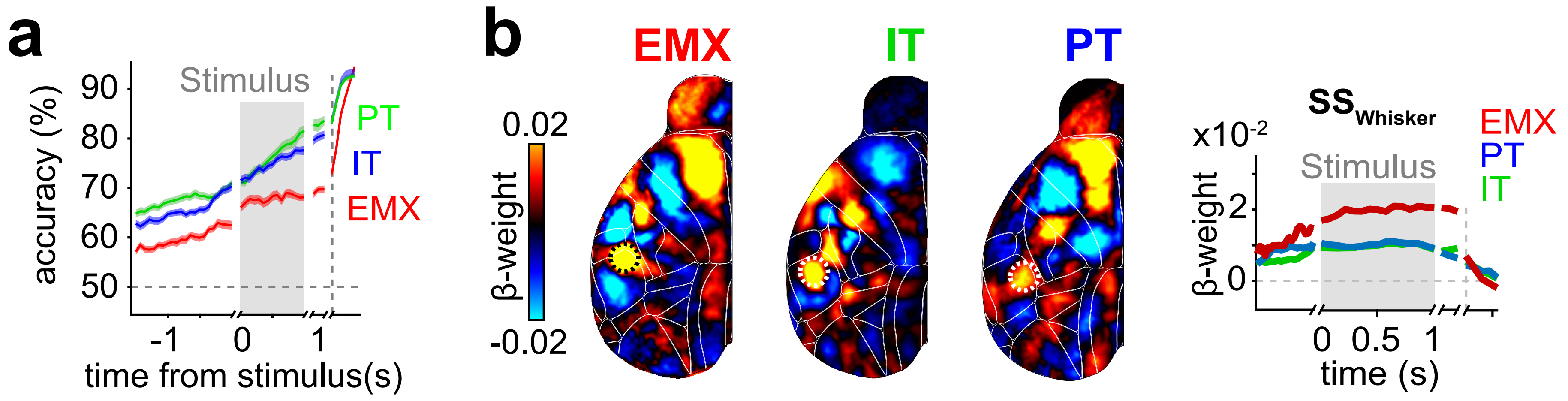


**Supplementary Fig. 10. Choice decoder performance and choice signals in somatosensory cortex**

**a)** 10x cross-validated decoder performance, predicting animal's left/right choices at different times during the trial. In all PyN types, decoder performance was above chance at all times, including the initiation period before the stimulus (gray box). This suggests that, in some trials, animals follow a pre-conceived choice that is stimulus-independent and can be decoded from cortical activity. Decoder performance was highest in the response period (dashed vertical line) when animals performed licking movements. **b)** Contralateral choice weight maps during the delay period (same as in Fig. 6c). Dashed circles show the location of somatosensory whisker cortex ($SS_{Whisker}$). In all PyN types, choice weights in $SS_{Whisker}$ were increased in the initiation period before the stimulus (gray box). A potential explanation could be that pre-stimulus choices are reflected in choice-specific whisker movements. However, choice signals in $SS_{Whisker}$ persisted when removing movement-related activity from the imaging data (Supp. Fig. 12). Whisker or other facial movements might therefore be too subtle to be captured by our analysis or choice signals in $SS_{Whisker}$ reflect non-overt choice-related activity.

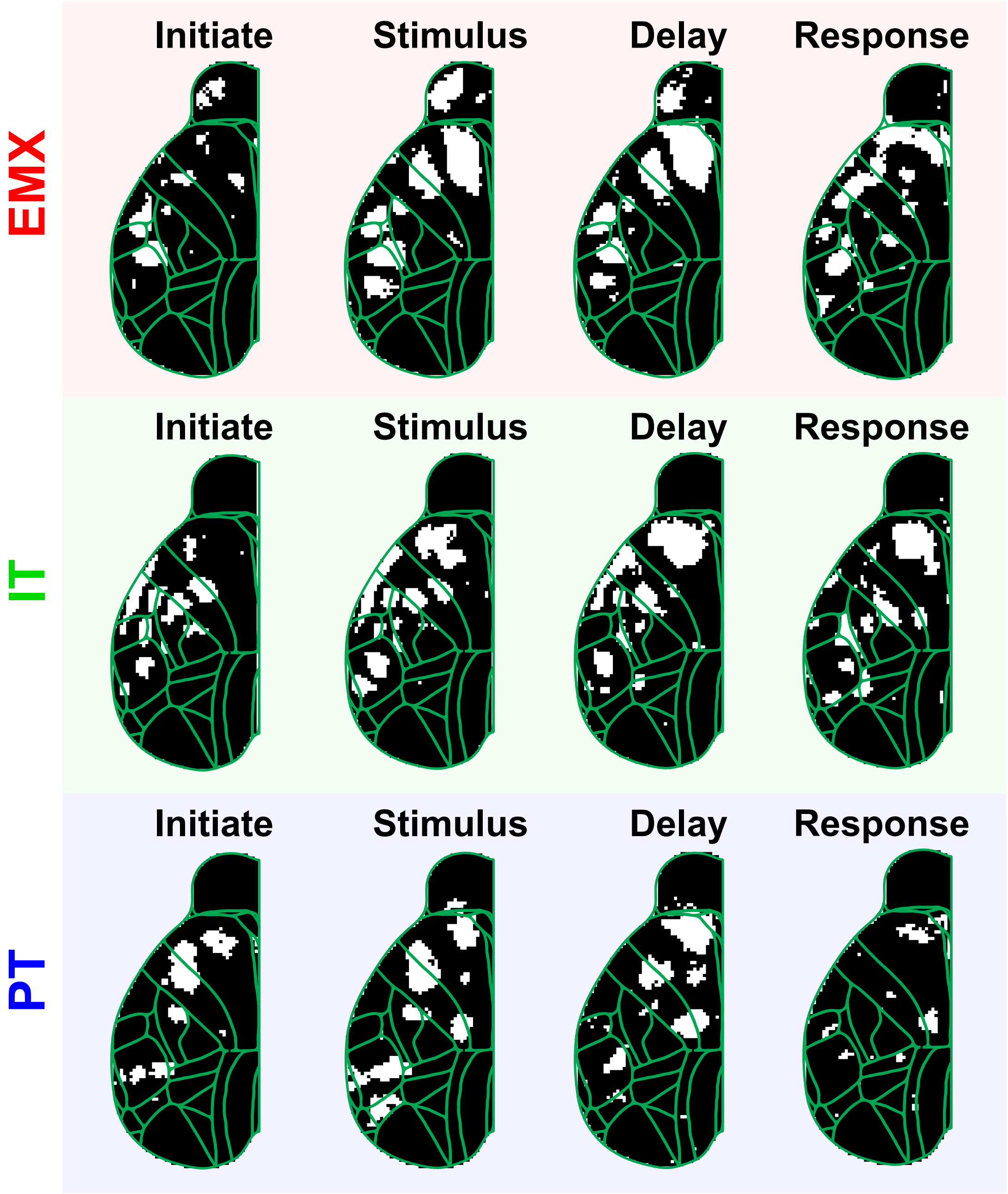


**Supplementary Fig. 11. Significance of choice decoder weights**

To assess significant weights of the choice decoder, we combined spatially downsampled choice maps from all sessions in each PyN type and subsequently performed a *t*-test in each pixel to determine which decoder weights are significantly different from zero. The resulting maps show significant pixels for different trial periods in white (*t*-test, $p < 0.05$, bonferroni-corrected for 3364 pixels). Significant pixels closely match choice decoder weights (Fig. 6c) with significant regions being largely tied to anterior cortex. In all PyN types, weights in frontal cortex are significant in the stimulus and delay period, thus supporting the main conclusions of the choice kernel analysis.

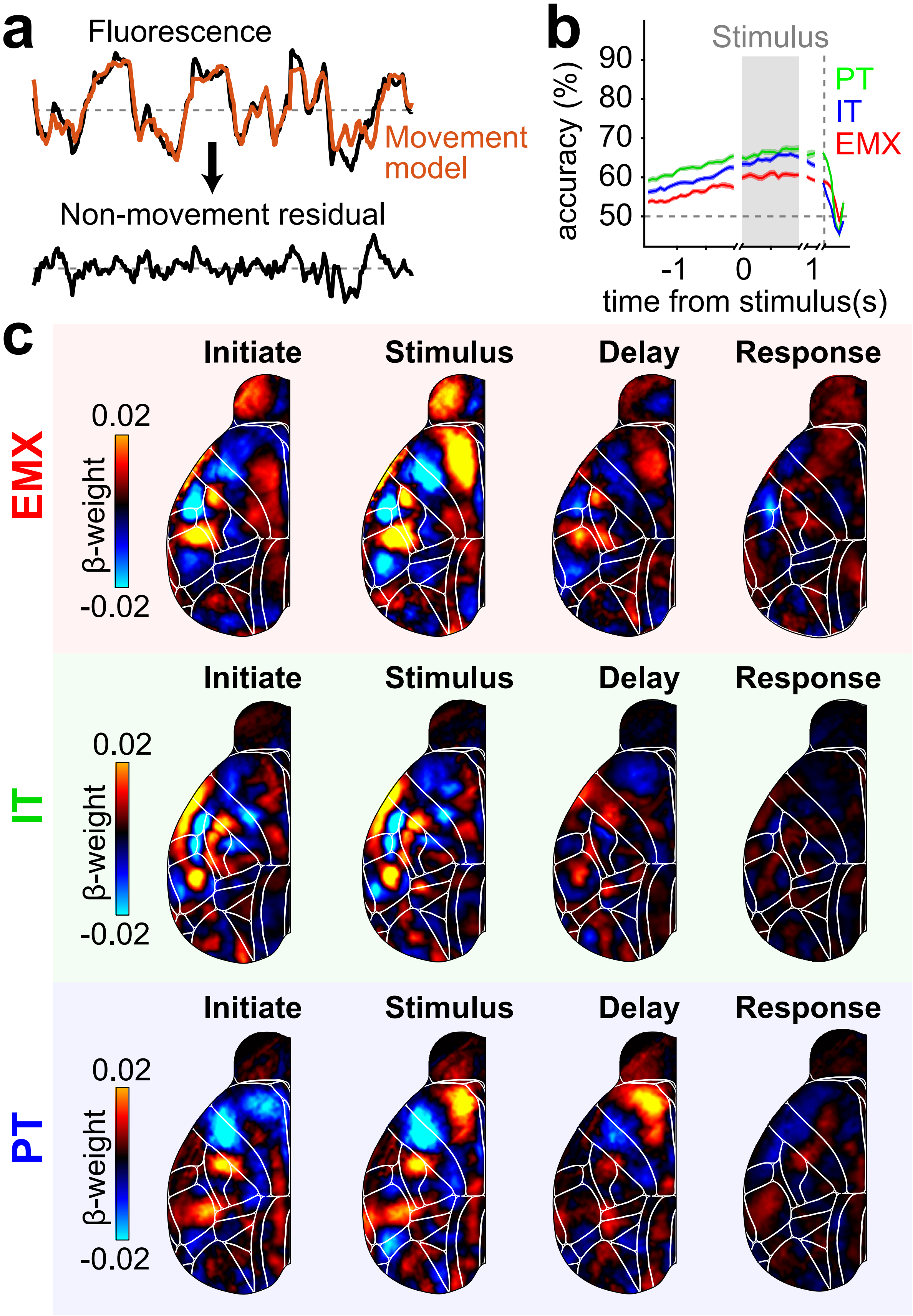


**Supplementary Fig. 12. Movement-corrected choice decoder**
**a)** Using a model based on movement variables (Supplementary Table 1), we subtracted all movement-related activity from raw fluoresence data and applied the choice decoder analysis to the resulting residuals. **b)** Removing movement-related activity reduced choice prediction accuracy, in particular during the delay and response period when most choice-related movements occur. In all PyN types, predictions remained above chance levels, suggesting that part of the choice-related activity is independent of observable movements. **c)** Movement-corrected choice kernels revealed the same cortical patterns as seen in the regular choice decoder (Fig. 6c), demonstrating that choice signals are not solely driven by movements.

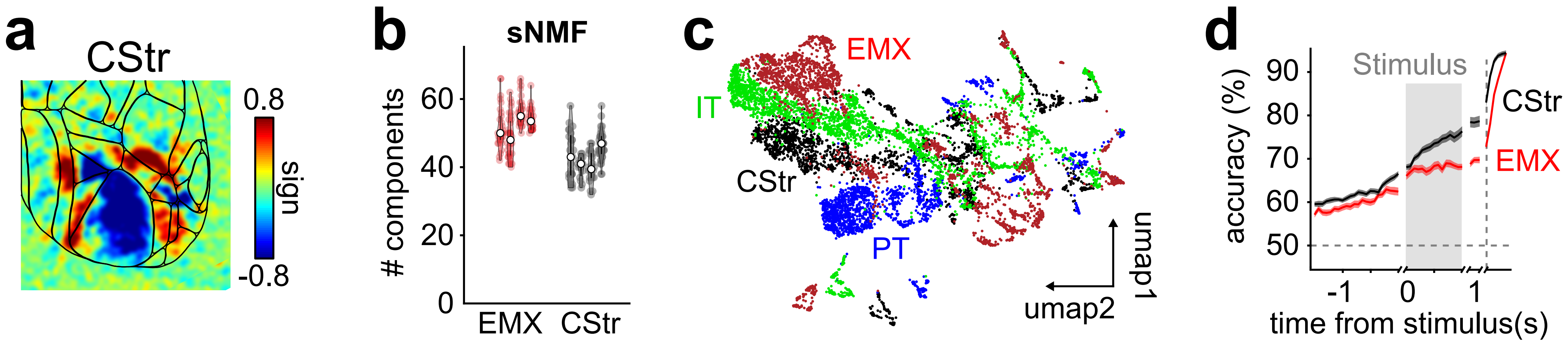


**Supplementary Fig. 13. Retrograde labeling of CStr neurons reveals distinct cortical dynamics**
**a)** Visual sign maps from retinotopic mapping experiments. CStr neurons responded to visual stimulation and reveal comparable retinotopic organization as other PyN types. **b)** Number of sNMF components, accounting for 99% of cortical variance in EMX and CStr mice, dots represent individual sessions. CStr neurons required less components as EMX and IT but more components as PT neurons (compare with Fig. 2a). **c)** UMAP embedding of spatial sNMF components for EMX (red), IT (green), PT (blue) and CSt (black) mice. Dots show individual spatial components. CStr components were clearly distinct from other PyN types. **d)** Cross-validated choice-decoder accuracy. Results are shown for EMX (red) and CStr mice (black). Decoder accuracy continuously increased throughout the trial for all PyN types. Dashed line indicates time of response, gray area is the stimulus period.

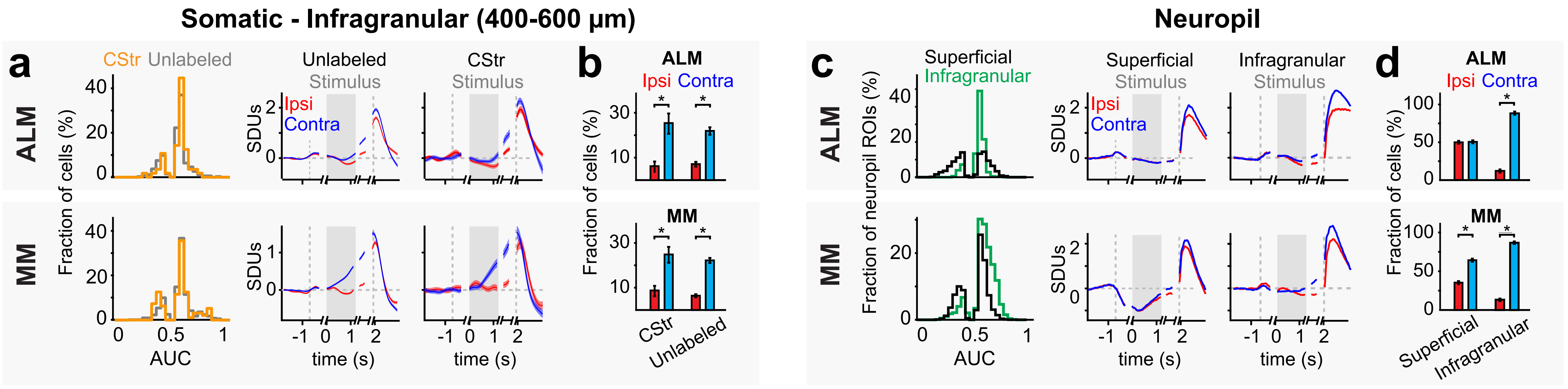


**Supplementary Fig. 14. Infragranular CStr neurons are tuned to contralateral choices**

**a)** Left: Overview of significantly choice-tuned neurons in deeper layers (400-600 µm) of ALM (top) and MM (bottom). Orange line: CStr neurons, labeled by tdTomato. Gray lines: unlabaled PyNs. AUC values below 0.5 indicate stronger responses for ipsilateral choices. Right: Trial-averaged activity for all choice-selective neurons, separated for ipsi- (red) versus contralateral choices (blue). Both CStr and unlabaled neurons show strong contralateral choice tuning with no clear difference between PyN-types. This suggest that ipsilateral choice tuning is limited to IT-CStr neurons in superficial layers of ALM. **b)** Fraction of cells responding selectively for ipsi- (red) versus contralateral choices (blue) in ALM and MM. CStr and unlabaled neurons in both show similar contralateral choice tuning. **c)** To differentiate somatic versus neuropil choice signals, we quantified ipsi- and contralateral choice tuning for neuropil ROIs. Each neuropil ROI represents the background fluorescence that sorrounded a given somatic ROI. Conventions as in a). **d)** Fraction of neuropil ROIs responding selectively for ipsi- versus contralteral choices. Conventions as in b). Neuropil ROIs were equally tuned to ipsi- and contralateral choices in superficial ALM layers but otherwise showed contralateral choice-specifity, generally recapitulating choice-specificity from unlabeled neurons. The symmetry found in neuropil choice tuning might explain the bilateral ALM activation observed with IT-specific widefield imaging (Fig. 3c), suggesting that IT-specific widefield signals are comprised of somatic and neuropil activity in the superficial cortex. In contrast, the stronger choice-selectivity in EMX- and PT-specific widefield imaging, suggests that these signals may emerge from infragranular neural activity .

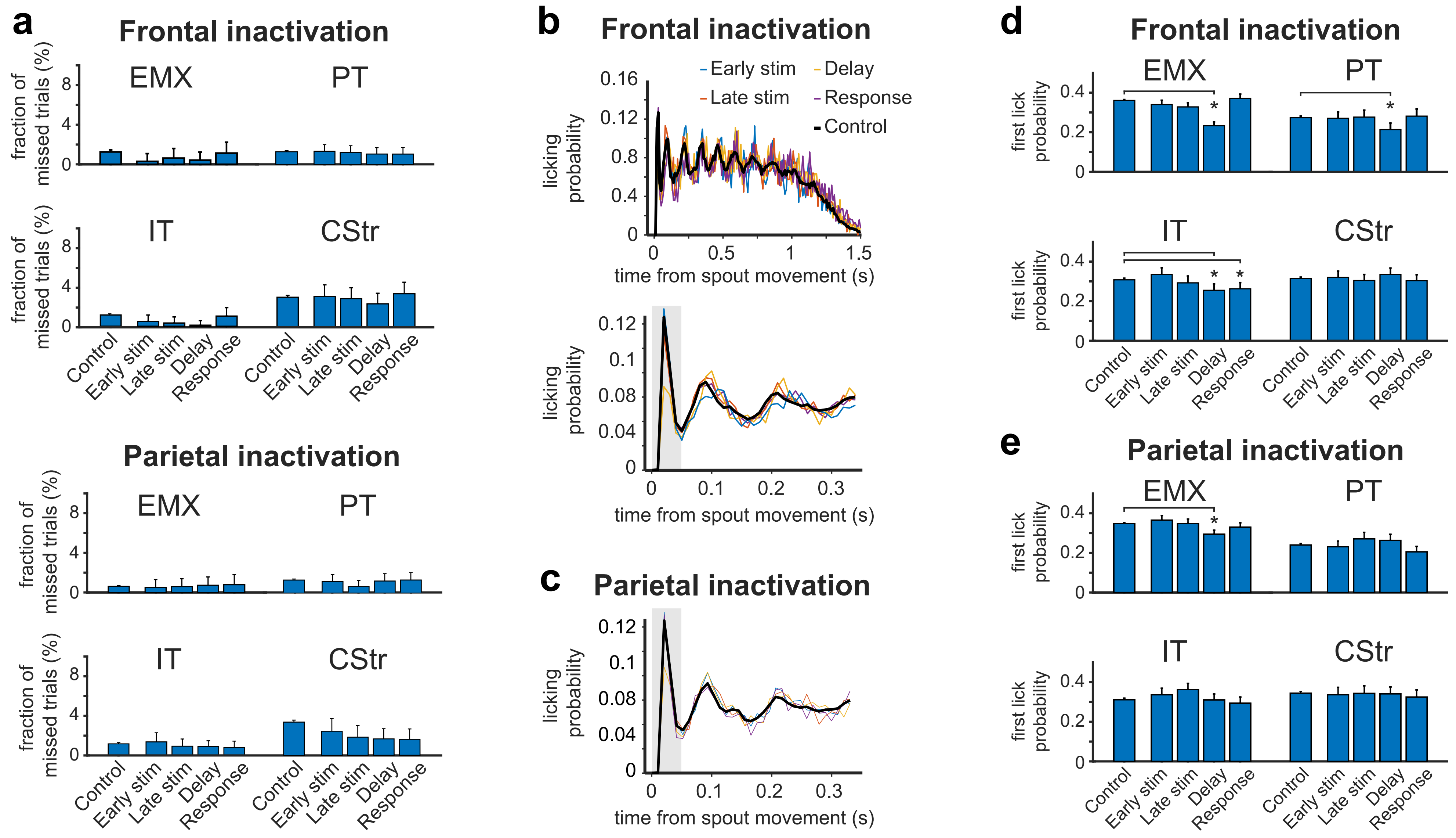


**Supplementary Fig. 15. Effects of optogenetic perturbation in frontal cortex on licking behavior**

**a)** To test if frontal or parietal inactivation impairs movement planning or execution, we computed the fraction of missed trials with and without optogenetic inhibition. No signficiant increase in missed trials was seen for frontal (top) or parietal inactivation (bottom), demonstrating that the animals' ability to respond was not impaired. **b)** Top: Quantification of licking behavior after spouts were moved in for all correct trials in a single EMX animal (10 ms bins). Licking probability varies rythmically at ~10 Hz as the animal licks the spout repeatedly (black line). The same pattern is observed with frontal optogenetic inactivation in different trial episodes (colored lines), demonstrating that motor generation is not generally perturbed. Bottom: While the lick pattern is largely similar with optogenetics, inactivation during the delay period (yellow line) reduces the lick probability during the first 40 ms (gray area). Frontal inactivation during the delay might thus increase animals' reaction times. **d)** Same as in b) but for parietal inactivation. **d)** Quantification of lick probability in the first 40 ms for all cell types. Frontal inactivation during the delay period reduces early lick probability in EMX, PT and IT mice but not CStr mice (stars indicate $p < 0.001$, binomial test). **e)** Same as in d) for parietal inactivation. Only EMX inactivation during the delay caused a small reduction in first lick probability.

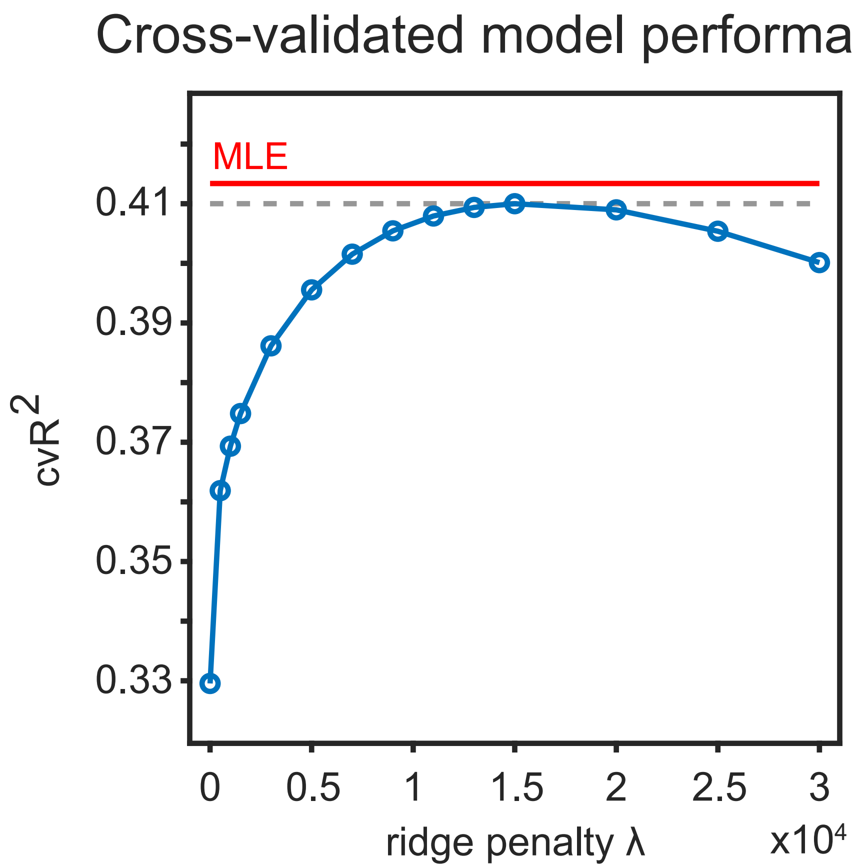


**Supplementary Fig. 16. Quantification of ridge penalty λ**
Quantification of different ridge λ penalties on cross-validated explained variance of the encoding model. λ is used to reguralize the model's predictor matrix and enforces sharing weights across correlated predictors. Increasing λ improves the cross-validated explained variance of the model, demonstrating that high λ values are beneficial to avoid overfitting.
Identification of the ideal λ through cross-validation (dashed gray line) is computationally expensive and required 38,6 minutes on a standard workstation PC for the example session above. We therefore used marginal maximum likelihood estimation (MLE), which allows to identify the ideal λ without cross-validation. This allows efficient identification of λ values for each of the 200 SVD components in the example recording (49,2 seconds in total), resulting in improved cross-validated reconstruction accuracy (solid red line).
